# Arkadia-SKI/SnoN signaling cascade differentially regulates TGF-β-induced iTreg and Th17 cell differentiation

**DOI:** 10.1101/2021.02.23.432519

**Authors:** Hao Xu, Lin Wu, Henry Nguyen, Kailin R. Mesa, Varsha Raghavan, Vasso Episkopou, Dan R. Littman

## Abstract

TGF-β signaling is fundamental for both Th17 and regulatory T cell (Treg) differentiation. However, these cells differ in requirements for downstream signaling components, such as various SMAD effectors, for their differentiation. To further characterize the mechanisms that distinguish TGF-β signaling requirements for Th17 and Treg cell differentiation, we investigated the role of Arkadia (RNF111), a RING type E3 ubiquitin ligase known to enhance TGF-β signaling during development. We find that Arkadia mediates the differentiation of induced-Treg (iTreg) but not Th17 cells both *in vitro* and *in vivo*. Inactivation of Arkadia in CD4^+^ T cells resulted in impairment of Treg cell differentiation *in vitro* and loss of RORγt^+^FOXP3^+^ iTreg cells in the intestinal lamina propria *in vivo*, which increased susceptibility to microbiota-induced mucosal inflammation. Furthermore, genetic ablation of two substrates of Arkadia, the transcriptional co-repressors SKI and SnoN, rescued *in vitro* and *in vivo* iTreg cell-differentiation of Arkadia-deficient cells. These results reveal distinct TGF-β signaling modules that govern Th17 and iTreg cell differentiation programs and that could be exploited therapeutically to selectively modulate T cell subsets in pathological settings such as autoimmunity or cancer.

## Introduction

T-helper 17 (Th17) and regulatory T cells (Treg) often act in opposition to each other in order to maintain balanced adaptive immune responses. Th17 cells are critical effector T cells in the maintenance of mucosal barrier homeostasis. However, uncontrolled Th17 responses can also contribute to autoimmune diseases such as spondyloarthropathies, psoriasis and inflammatory bowel disease in both mouse and human^1^. Treg cells modulate the immune system to maintain tolerance to both self- and nonself-antigens, thus preventing autoimmune diseases and restraining excessive tissue inflammation. Conversely, unchecked Treg cell activity can lead to suppression of critical immune functions such as tissue immunosurveillance and tumor cell elimination^2, 3^.

Cytokine-activated STAT3 and transforming growth factor–β (TGF-β) signaling pathways are vital for Th17 cell differentiation^4, 5, 6, 7, 8, 9^, while signaling through the IL-23 and IL-1 receptors is additionally required for Th17 cell-mediated pathogenicity in mouse auto-immune disease models^10, 11, 12, 13^. Under certain conditions however, TGF-β is arguably dispensable for Th17 cell differentiation. For instance, in the absence of TGF-β, IL-6 alone can induce the Th17-regulating transcription factor RORγt and IL-17A/F expression in STAT6/T-bet double knockout T cells both *in vivo* and *in vitro*^14, 15^. Taken together with the reciprocal suppressive effects of IL-2/STAT5 on RORγt expression, and IL-6/STAT3 towards the Treg transcription factor Foxp3^5, 16^, these data are consistent with key roles of inhibitory pathways in restraining alternative differentiation programs. Nevertheless, TGF-β signaling is required for Th17 responses in physiological conditions. In TGF-β1 knockout mice, IL-17A-producing T cells were diminished or absent in spleen, mesenteric lymph nodes and lamina propria, and there was less IL-17A in serum^9^. Furthermore, mice failed to develop Th17-mediated experimental allergic encephalomyelitis (EAE) when a dominant negative form of TGF-β receptor II (TGFBRII) was expressed under control of the *Cd4* promoter (*Cd4dnTgfbr2*)^17^. Similarly, there were reduced proportions of IL-17A^+^ or RORγt-expressing T cells in the intestinal lamina propria of *Cd4dnTgfbr2* mice or in mice with TGF-β receptor I deficiency in T cells^13^.

There are two subsets of Treg cells, thymus-derived natural Treg (nTreg), constituting a central arm of immunological tolerance to self-antigens, and peripherally-induced Treg cells (iTreg) arising from recognition of non-self-antigens. TGF-β signaling is important for the differentiation of both subsets of Treg cells. In neonatal mice lacking *Tgfbr2* in T cells^18,19^, nTreg cells were reduced due to excessive negative selection of CD4^+^CD8^+^ double positive and CD4^+^ single positive thymocytes. The loss of nTreg cells was transient and restored quickly in two weeks, which was attributed to enhanced IL-2 expression in the thymus^18^. Differentiation of iTreg cells relies on both the TGF-β signaling cascade^20, 21^ and the IL-2/STAT5 signaling pathway^22, 23^. There was substantial decrease in iTreg cells in lamina propria of mice with T cell-restricted deficiency of *Tgfbr1*^13^. Consistent with this finding, there was impaired commitment of peripheral iTreg cells in mice with targeted deletion of the *Foxp3 CNS1* enhancer^24, 25^, which contains binding sites for the SMAD2/3 transcriptional effectors, which act downstream in the TGF-β signaling cascade^26^. Together these studies highlight the importance of TGF-β in the differentiation of both Th17 and Treg cells.

While both Th17 and iTreg cell differentiation rely on signaling through TGF-β receptors *in vivo*, deficiencies of downstream signaling components show divergent effects on the differentiation programs. TGF-β binds to a hetero-tetrameric receptor complex composed of TβRI and TβRII and triggers the canonical SMAD pathway as well as other signaling modalities, including MAPK and PI3K pathways. In the canonical SMAD pathway, SMAD2 or SMAD3 (SMAD2/3), phosphorylated by TβRI, form a complex with SMAD4 and translocate into the nucleus for transcriptional regulation (Reviewed in ^27^). SMAD4 knockout (KO) or SMAD2/3 double knockout (DKO) T cells lose their ability to induce *Foxp3* and differentiate into iTregs^28, 29^. However, Th17 cell differentiation in the SMAD4 KO background remained similar to that of wild type mice both *in vitro* and *in vivo*^28^. Furthermore, in SMAD2/3 DKO cells the amount of RORγt was similar to wild type levels, although there was reduced IL-17A upon stimulation with TGF-β plus IL-6^29^. The proposed explanation was that the complex between SMAD4 and the corepressor SKI inhibits the expression of *Rorc(t)*, which encodes RORγt. Thus, knockout of SMAD4 or TGF-β stimulation, resulting in degradation of SKI, would lead to the removal of inhibition of *Rorc(t)* expression and promote the Th17 program^30^. Indeed, inactivation of SMAD4 was sufficient to permit induction of RORγt expression by IL-6 in the absence of TGF-β^30^. In contrast to their function in Th17 cells, the SMAD complexes act as transcriptional activators to induce *Foxp3* expression in iTreg cells, but the differences in mechanisms between the cell types remain unclear.

We re-examined TGF-β signaling in Th17 and iTreg cell differentiation, and found that SMAD4 deficiency had a limited effect on the induction of Th17 cells, but resulted in loss of iTreg cell differentiation both *in vitro* and *in vivo*. To further explore potential differences between Th17 and iTreg cells in the TGF-β signaling pathway, we examined the role of Arkadia (RNF111), an E3 ligase involved in multiple signaling pathways mediated by TGF-β family members. Arkadia promotes NODAL signaling during mesendoderm specification and establishment of the body axis, targeting the degradation of the transcriptional co-repressors SKI and SnoN, as well as members of the SMAD family ^31, 32, 33, 34, 35^. We found that Arkadia was selectively required for iTreg, but not Th17, cell differentiation, and its absence in T cells resulted in increased susceptibility to inflammatory bowel disease. Absence of Arkadia resulted in failure of TGF-β-induced SKI and SnoN degradation, and targeted inactivation of these co-repressors rescued Foxp3 induction in Arkadia-deficient T cells. Our results thus identify another key distinction in TGF-β signaling pathways in Th17 and iTreg cells, which could potentially be exploited for selective therapeutic targeting of these T cell subsets.

## Results

### Arkadia regulates *in vitro* differentiation of Treg but not Th17 cells in response to TGF-β

We examined *in vitro* Th17 cell differentiation with cells from mice with T cell-selective inactivation of *Smad4*. Consistent with published data^30^, loss of *Smad4* had little effect on the induction of RORγt, which was observed under conventional Th17 cell differentiation conditions with TGF-b and IL-6 and even when only IL-6 was present (Extended Data Fig. 1a). However, in contrast to the earlier study, there was little expression of IL-17A in SMAD4-deficient cells after exposure to IL-6. Consistent with the importance of TGF-β signaling in Th17 cell differentiation, there was marked reduction in the proportion of RORγt^+^FOXP3^-^ Th17 cells in the large intestine lamina propria (LILP) of mice with T cell-specific inactivation of both SMAD4 and TGFβR2 (Extended Data Fig. 1b, c). We next used CRISPR/Cas9 gene-targeting to interrogate the roles of multiple TGF-β signaling pathway genes in regulating Th17 and iTreg cell differentiation *in vitro*. As expected, inactivation of *Smad4* or *Stat5* inhibited the differentiation of iTreg but not Th17 cells (Extended Data Fig. 2a). Among additional targeted genes, we were particularly interested in examining the role of Arkadia/RNF111, which was shown to enhance the TGF-β-SMAD2/3 signaling pathway^31, 32, 33, 36^.

Arkadia was first identified as an essential component of a subset of NODAL signaling responses and functions during embryonic development^37, 38^. Subsequent studies showed that TGF-β signaling activates Arkadia to unibiquitinate and degrade SKI and SnoN. SKI and SnoN bind to SMAD4 and ligand-activated SMAD2/3 and recruit transcriptional co-repressor complexes containing histone deacetylases (HDAC) to suppress TGF-β-induced transcription^39, 40, 41^. Because TGF-β signaling was proposed to relieve Smad4- and SKI-dependent inhibition of *Rorc(t)* expression and Th17 cell differentiation^30^, we hypothesized that Arkadia may mediate the TGF-β-induced degradation of the SMAD4-associated repressive complex. In contrast to our expectation, inactivation of Arkadia had no effect on the induction of RORγt under Th17 cell differentiation conditions (Extended Data Fig. 2a). Instead, there was dramatic loss of FOXP3 under Treg differentiation conditions. Similar results were observed with multiple guide-RNAs (gRNAs) targeting diverse regions of the *Arkadia* gene (Extended Data Table I and data not shown).

Since global knockout of Arkadia in mice results in embryonic lethality, we generated *Arkadia^fl/fl^* mice and bred them to *Cd4^Cre^* transgenic mice to investigate the role of Arkadia in T cell biology. *Arkadia^fl/fl^ Cd4^Cre^* mice were viable, fertile and normal in size and did not display obvious signs of inflammation. Draining lymph nodes and spleen displayed similar cellularity in both conditional knockout mice and control littermates. There were no differences in proportions of CD44^-^CD62L^+^CD4^+^ and CD44^+^CD62L^-^CD4^+^ cells in spleen (data not shown). Arkadia-deficient T cells expressed levels of IFN-γ, IL-17A and RORγt similar to control cells under corresponding *in vitro* differentiation conditions. In contrast, there was substantial reduction of Foxp3 expression in Arkadia-deficient T cells under Treg differentiation conditions (Fig. 1a-c). To confirm that Arkadia function is required for Treg cell differentiation *in vitro*, exogenous wild type or E3 ligase activity-dead Arkadia proteins were expressed in Arkadia-deficient T cells. Wild type, but not the mutant Akadia protein, rescued *Foxp3* expression in T cells differentiated under Treg conditions (Fig. 1d, e). Under suboptimal conditions for Th17 cell differentiation, achieved at higher concentrations of TGF-β, T cells deficient for Arkadia retained their ability to produce IL-17A. By contrast, the proportions of IL-17A-producing cells were reduced in wild type cells with increasing concentration of TGF-β, due to upregulation of *Foxp3*, as we previously showed (Extended Data Fig. 2b)^42^. Collectively, these results indicate that Arkadia is required for induction of *Foxp3* expression in response to TGF-β.

**Figure 1.**
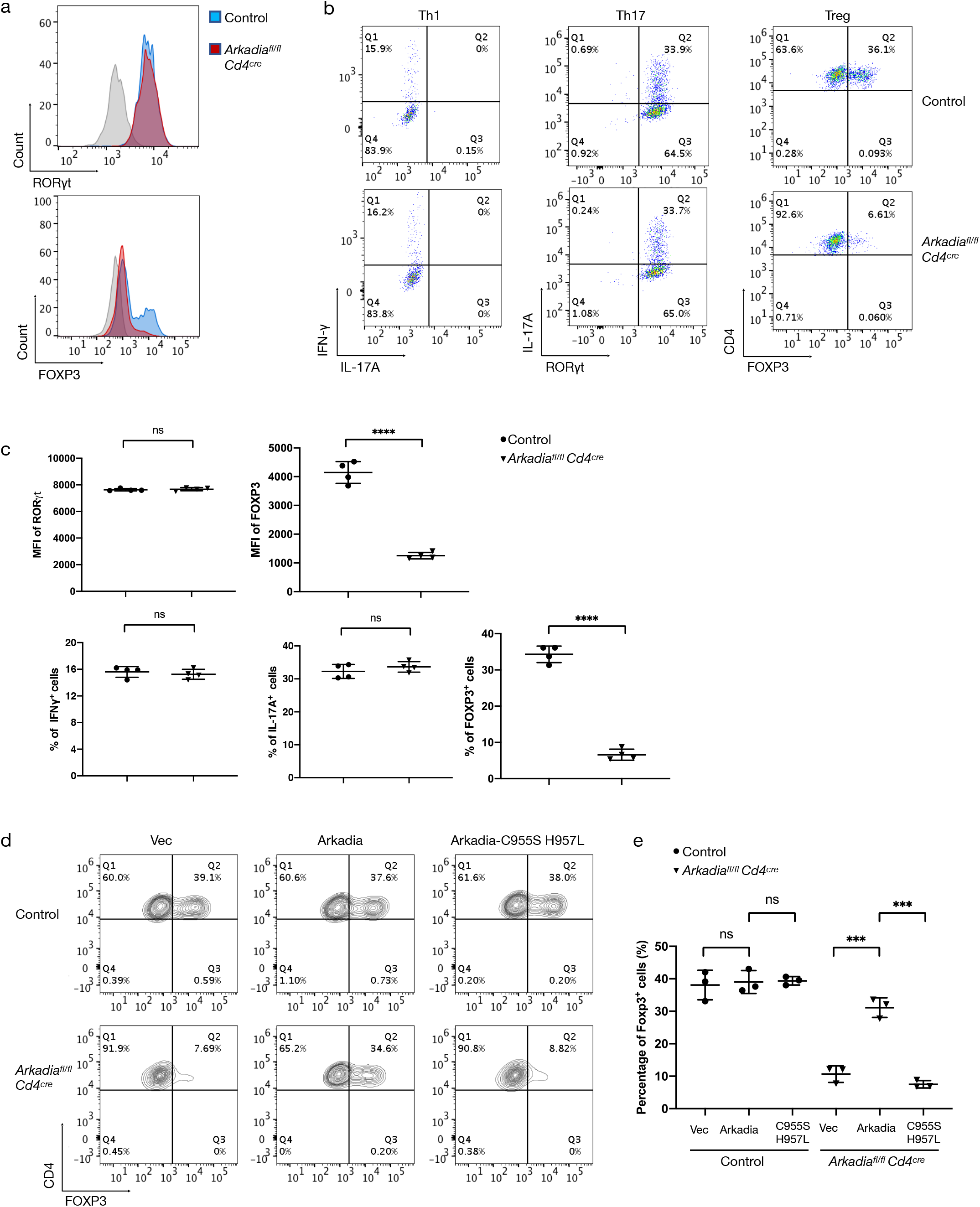
Arkadia is selectively required for in vitro Treg cell differentiation. (a) Expression of RORγt and FOXP3 in WT and *Cd4^Cre^ Arkadia^fl/fl^* T cells differentiated *in vitro* under Th17 and Treg conditions, respectively. (b) Representative flow cytometry profiles of IFN-γ, IL-17A, RORγt and FOXP3 in T cells from WT and mutant mice following *in vitro* differentiation under Th1, Th17 and Treg conditions. (c) Statistical analysis of MFI (Mean Fluorescence Intensity) of RORγt and FOXP3 (top panels) and percentage of cells expressing IFN-γ, IL-17A, and FOXP3 (bottom panels) following differentiation as in (a, b). (d, e) Rescue of Treg cell differentiation in *Arkadia*-deficient CD4^+^ T cells. Representative flow cytometry panels (d) and composite data (e) for FOXP3 expression in control and *Arkadia* knockout Treg cells transduced with lentivirus expressing empty vector (Vec), wild type Arkadia (Arkadia), and Arkadia E3 ligase activity-dead mutant proteins (Arkadia-C955S H957L). Data (a-e) are representative of three independent experiments. Control, CD4^+^ cells from control mice; *Arkadia^fl/fl^ Cd4^cre^*, CD4^+^ cells from *Arkadia* knockout mice. Statistics were calculated using unpaired t test. ns = not significant, *** *P* < 0.001, and **** *P* < 0.0001.

### Requirement for Arkadia in iTreg cell differentiation *in vivo*

To further explore the role of Arkadia in the genesis of Treg cells, we analyzed the development of thymus-derived nTreg cells and the differentiation of peripheral iTreg cells in Arkadia conditional mutant mice and control littermates. *Arkadia^fl/fl^ Cd4^Cre^* mice had normal numbers of developing thymocyte subsets, but displayed an increase in the proportion and number of FOXP3^+^Helios^+^ cells among CD4 single positive cells (Extended Data Fig 3a, b). This finding might be due in part to enhanced IL-2 expression that accompanies disrupted TGF-β signaling in the thymus^18^. Consistent with impaired Treg cell polarization *in vitro*, RORγt^+^FOXP3^+^ iTreg cells were markedly reduced in the LILP of Arkadia conditional mutant mice. Interestingly, RORγt^+^FOXP3^-^ Th17 cells were increased in both proportion and absolute cell number when Arkadia was absent in T cells (Fig. 2a). There were no differences in GATA3^+^ cells among the FOXP3^+^ and FOXP3^-^ populations, or in IFNγ and IL-17A production among LILP CD4+ T cells from WT and *Arkadia^fl/fl^ Cd4^Cre^* mice (Extended Data Fig. 3c, d). Together, these results suggest that Arkadia specifically regulates iTreg cell differentiation *in vivo*.

**Figure 2.**
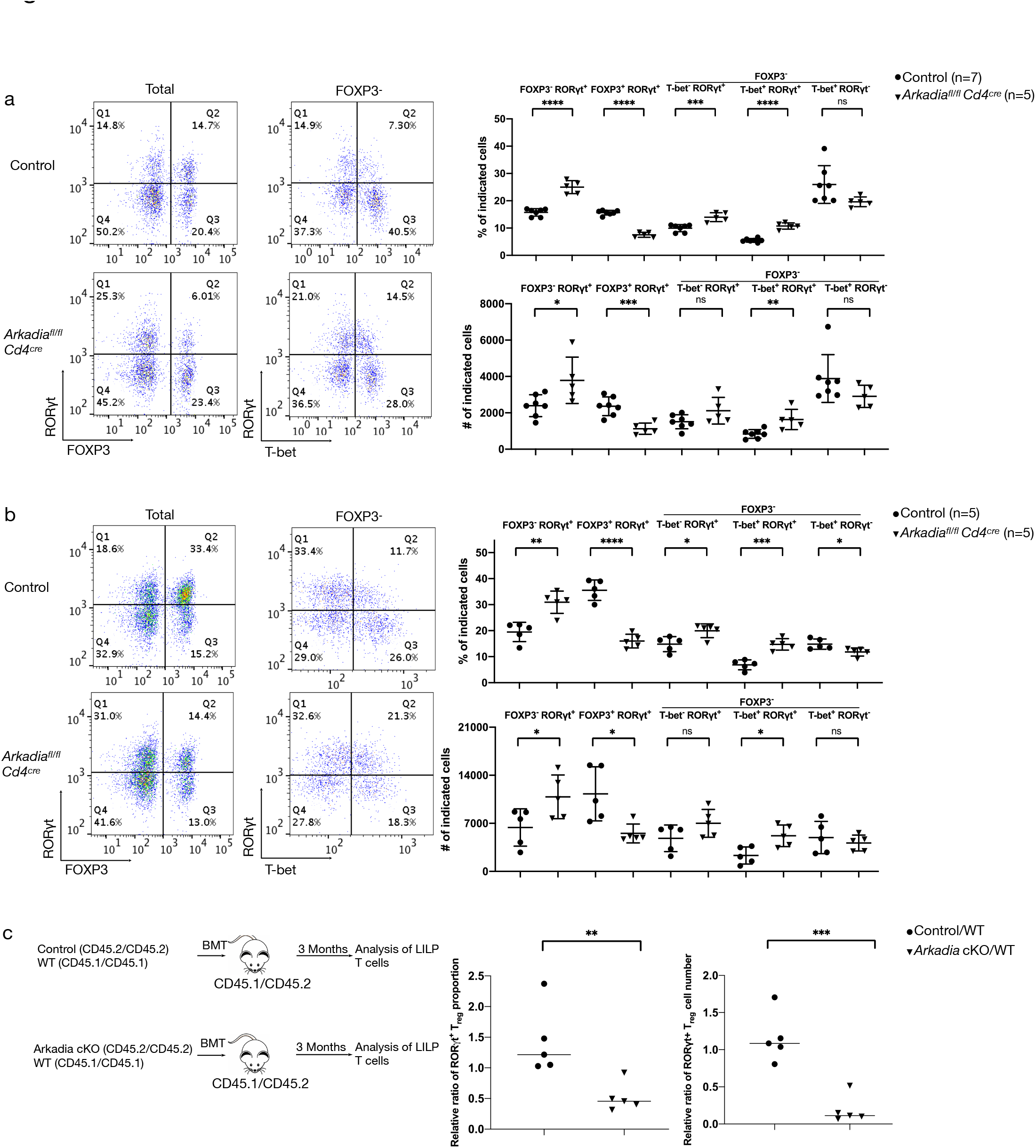
Arkadia is required for iTreg but not Th17 cell differentiation *in vivo*. (a) CD4^+^ TCRβ^+^ T lymphocytes from LILP of control and *Arkadia* conditional knockout mice under SPF conditions were stained for RORγt, FOXP3 and T-bet. Left, representative flow cytometry plots; right, cell proportions and numbers of indicated T lymphocyte subpopulations, from 1 of 3 independent experiments. n = 13 in the 3 experiments. (b) Staining of RORγt, FOXP3 and T-bet in CD4^+^ TCRβ^+^ T lymphocytes from LILP of control and *Arkadia* conditional knockout mice colonized with *H. helicobacter*. Panels as in (a). Data are representative of 2 independent experiments (n = 10). (a, b) Statistical analysis were performed with unpaired t test. ns = not significant, * *P* < 0.05, ** *P* < 0.01, *** *P* < 0.001, and **** *P* < 0.0001. (c) Left panel, experimental scheme for mixed bone marrow chimera experiment. Bone marrow cells from control or *Arkadia* conditional knockout mice (CD45.2/CD45.2) were mixed with an equal number of bone marrow cells of CD45.1/CD45.1 mice (WT) and transferred into lethally irradiated CD45.1/CD45.2 recipient mice. CD4^+^ TCRβ^+^ T lymphocytes from LILP were stained for RORγt and FOXP3 at 3 months after transfer. Right panels, relative ratios of RORγt^+^ FOXP3^+^ lymphocytes in the reconstituted mice, based on their proportions among total CD4^+^ T cells and on their total numbers (designated as Control/WT and *Arkadia* cKO/WT, based on the isotype markers). The relative ratios were normalized to the ratios of donor B cells in peripheral blood. Ratios of co-transferred cells in each animal were calculated individually and combined for analysis with unpaired t test. ** *P* < 0.01 and *** *P* < 0.001.

*Helicobacter hepaticus* is a commensal bacterium that induces RORγt^+^ Treg cells to maintain intestinal homeostasis in wild type mice^43^. Mice were colonized with *H. hepaticus* to interrogate whether Arkadia deficiency in T cells would affect *H. hepaticus*-specific iTreg cell differentiation. As expected, the proportion and number of RORγt^+^FOXP3^+^ regulatory cells were substantially higher in *H. hepaticus*-colonized compared to uncolonized mice (Fig. 2a and 2b), and this increase was not observed in *Arkadia^fl/fl^ Cd4^Cre^* mice. Although the colonized mutant mice had higher numbers of RORγt^+^FOXP3^-^ Th17 cells, severe spontaneous colitis was not detected (Fig. 2b and Extended Data Fig. 3e). To confirm that the Arkadia-deficient T cells had a cell-intrinsic defect in iTreg cell differentiation, we performed mixed bone marrow chimera experiments. Bone marrow from control (wild type) or *Arkadia* conditional mutant mice (CD45.2/CD45.2) was mixed with bone marrow of CD45.1/CD45.1 mice (WT), followed by transfer into lethally irradiated CD45.1/CD45.2 recipient mice. Three months after transfer, analysis of LILP T lymphocytes revealed a reduction in the proportion and cell number of iTreg cells derived from Arkadia-deficient bone marrow, confirming that Arkadia is cell-intrinsically required for *in vivo* differentiation of iTreg cells, and not in another T cell subset that influences iTreg cell differentiation (Fig. 2c and Extended Data Fig. 4).

### Arkadia is required to maintain intestinal homeostasis

It is well-established that RORγt^+^ iTreg cells are critical in the maintenance of gastrointestinal homeostasis. Mice that completely lack these regulatory T cells develop minor to pronounced signs of pathology at mucosal sites with various inflammatory signatures, many of which are microbiota-dependent. Furthermore, it has been shown that mice with partially compromised iTreg cell differentiation, such as mutants for *CNS1* of *Foxp3, Cmaf* or *Tgfbr1* in Treg cells, which show no sign of pathology in young or immunocompetent mice, eventually develop intestinal inflammation and pathogenesis with age or immunodeficiencies^24, 25, 43, 44^. Therefore, we reasoned that mice with T cell-specific Arkadia deficiency might be more susceptible under conditions that sensitize for inflammation.

To test this hypothesis, we administered anti-IL-10RA antibody to mutant and control mice kept under SPF conditions with or without *H. hepaticus* colonization (Fig. 3a and Extended Data Fig. 5a). In mice with *H. hepaticus*, injection of 0.25 mg antibody was well-tolerated by control mice, but Arkadia conditional mutant mice developed severe colitis (Fig. 3a). Strikingly, *H. hepaticus* induced expansion of RORγt^+^ iTreg cells in control littermates, but proinflammatory Th17 cells with Th1-like features in *Arkadia^fl/fl^ Cd4^Cre^* mice. These proinflammatory Th17/Th1 cells were characterized by expression of both RORγt and T-bet, which was accompanied by marked reduction in RORγt^+^ FOXP3^+^ cells in the LILP (Fig. 3b and 3c). As expected, the proinflammatory Th1 or Th1-like Th17 cells expressed high levels of IFN-γ or both IL-17A and IFN-γ upon *ex-vivo* re-stimulation (Fig. 3d). Under SPF conditions, in the absence of *H. hepaticus*, there are fewer microbiota-induced iTreg cells and, upon blockade, fewer Th17/Th1 cells^43^. Accordingly, there was little intestinal inflammation and pathology in WT mice even when a higher dose (1 mg) of anti-IL-10RA antibody was administered. However, Arkadia mutant mice developed colitis under these conditions (Fig. 3e). Differences in LILP T cells between control and mutant mice were similar to those observed with the *H. hepaticus* colonization conditions (Extended Data Fig. 5a-d). The expansion of Th1 and Th1-like Th17 cells rather than iTreg cells in mice that developed inflammation in these colitis models further supports a specific requirement for Arkadia for iTreg cell differentiation. However, Th17 cell differentiation induced by segmented filamentous bacteria (SFB)^45^ was similar in Arkadia conditional knockout mice and control littermates. Furthermore, there was no significant difference in disease severity of Th17-mediated EAE between mice with Arkadia-deficient or -sufficient T cells, supporting our conclusion that Arkadia is dispensable for Th17 cell differentiation *in vivo* (Extended Data Fig. 6a, b).

**Figure 3.**
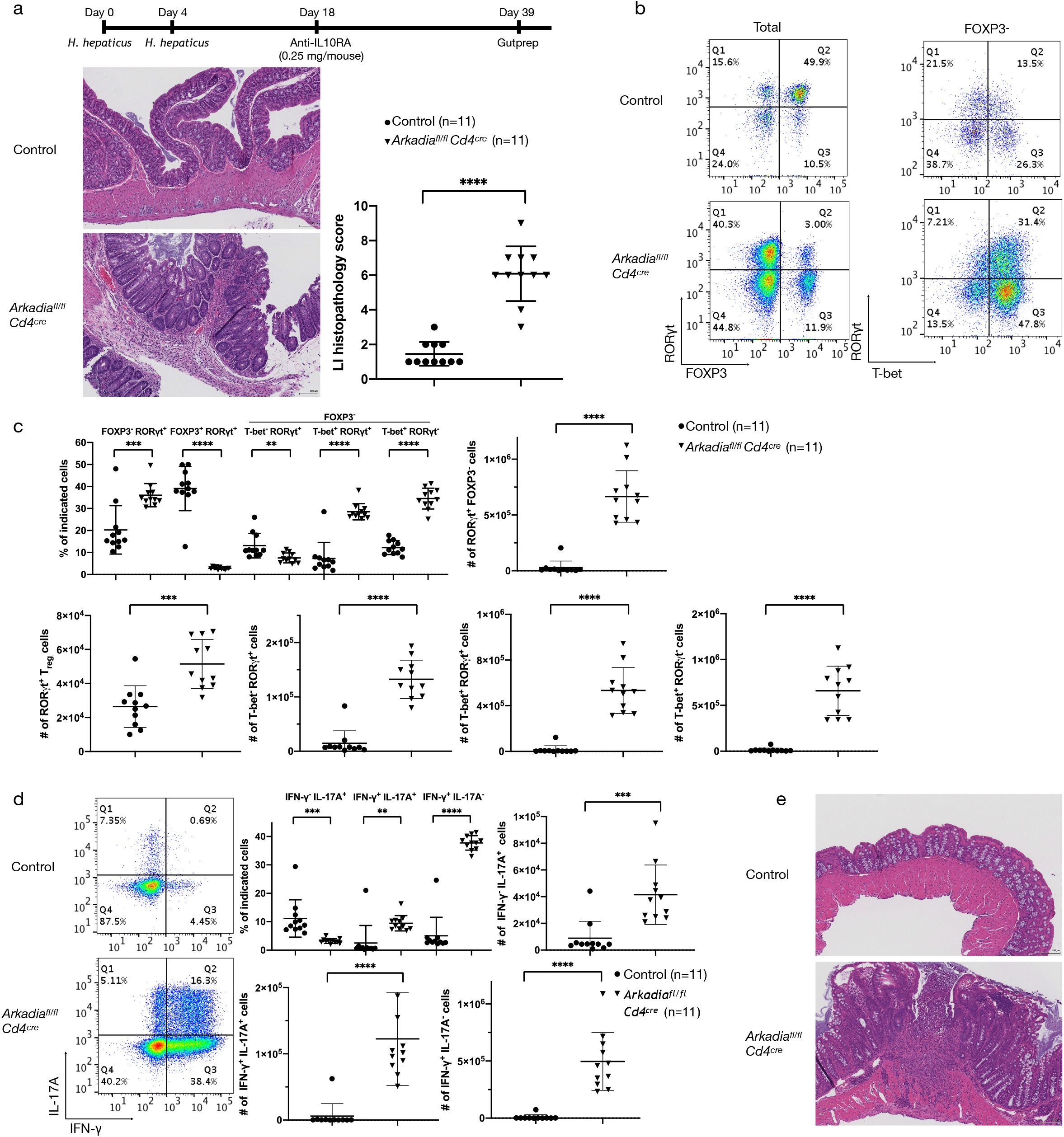
Arkadia is required in T cells to maintain mucosal homeostasis. (a) Upper panel, schematic of IL-10RA blockade in *H. hepaticus*-colonized mice. Lower panels, representative large intestine sections (left) and colitis scores (right). Data from two independent experiments (n=11) are combined. Statistical analysis was performed with unpaired t test. ns = not significant, **** *P* < 0.0001. (b) Representative RORγt, FOXP3 and T-bet staining in CD4^+^ TCRβ^+^ T lymphocytes isolated from LILP of indicated mice with IL-10RA blockade. (c, d) Flow cytometry and statistical analysis (cell proportion and number) of indicated T lymphocyte subpopulations with IL-10RA blockade shown in (b). c, statistical analysis of LILP CD4^+^ TCRβ^+^ lymphocytes expressing RORγt, FOXP3 and T-bet. d, representative flow cytometry analysis of LILP lymphocytes expressing INF-γ and IL-17A effector cytokines (left) and statistical analysis (cell proportion and number) of indicated subpopulations (right). Differences between experimental groups were determined by unpaired t test, ** *P* < 0.01, *** *P* < 0.001, and **** *P* < 0.0001. (e) Representative H&E staining of large intestine sections in mice with IL-10RA blockade maintained under SPF conditions (n=8).

### Arkadia degrades SKI/SnoN proteins to promote Treg cell differentiation

We next explored the mechanism by which Arkadia participates in iTreg cell differentiation, and focused on the transcriptional co-repressors SKI and SnoN, which Arkadia targets for degradation in response to TGF-β signaling^31, 36^. Our mRNA-Seq data showed that only *Ski* and *SnoN*, among 8 *Ski* family members, are highly expressed in *in vitro* differentiated Th17 and Treg cells (Extended Data Table II). We therefore assessed the degradation of SKI and SnoN in Arkadia-deficient or -sufficient T cells following TGF-β stimulation. Since current commercial antibodies recognize other proteins non-specifically, we used *Ski* and *SnoN* knockout mouse embryonic fibroblasts to confirm their specificity (Extended Data Fig. 7a). The turnover of SKI and SnoN was then investigated in differentiating Th17 and Treg cells. In response to TGF-β treatment, there was rapid degradation of both SKI and SnoN under Th17 and Treg differentiating conditions. However, both proteins remained intact in Arkadia-deficient T cells under the same conditions, consistent with their being targeted for degradation by the E3 ligase (Fig. 4a).

**Figure 4.**
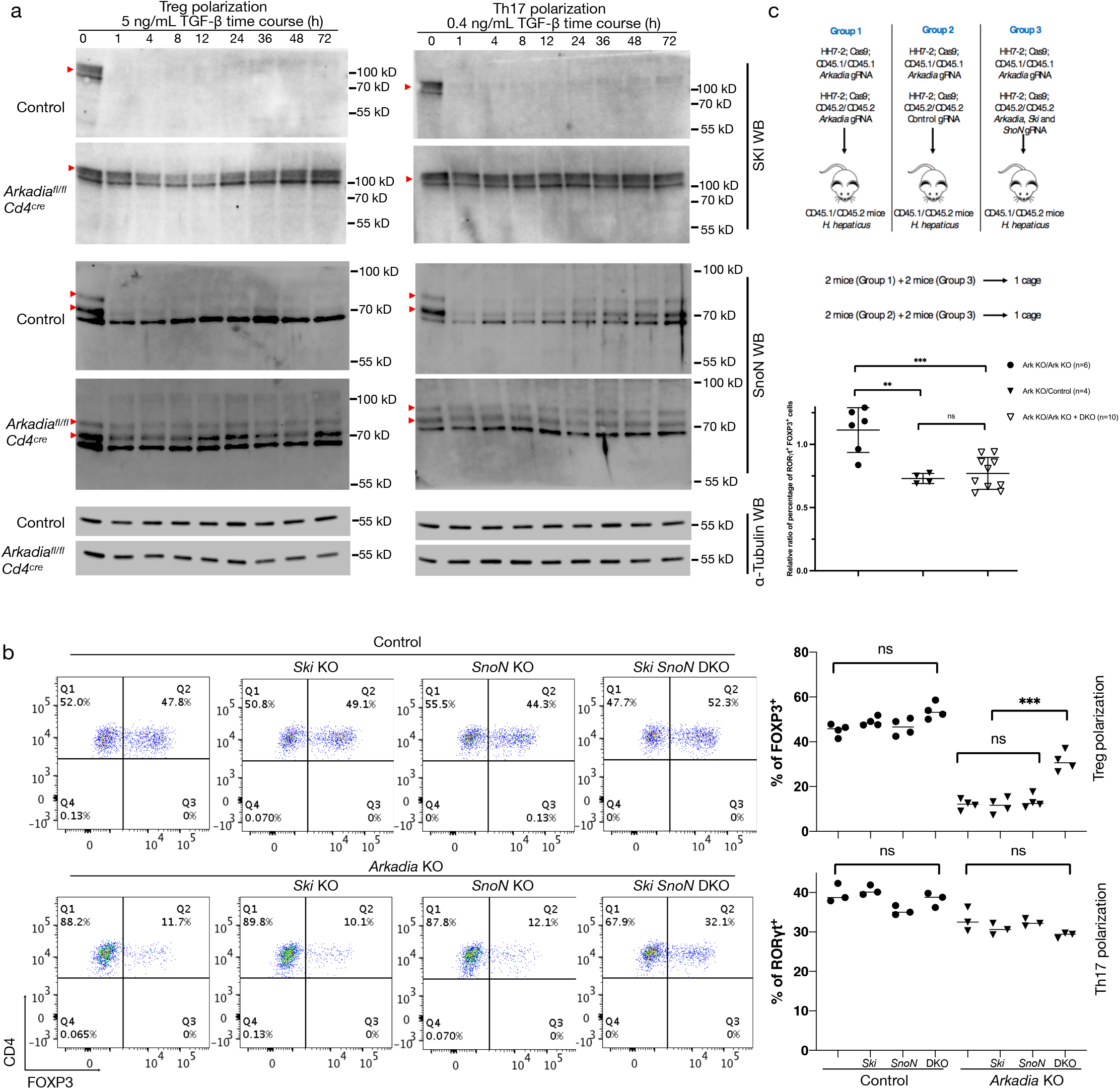
Arkadia regulates iTreg cell differentiation through degradation of SKI and SnoN. (a) Time course of degradation of SKI and SnoN proteins under *in vitro* Th17 and Treg differentiation conditions. Control or *Arkadia*-deficient T cells were polarized under Th17 or Treg conditions for indicated times, followed by SKI and SnoN immunoblotting analysis of cell lysates. Red arrowheads indicate SKI and SnoN proteins detected by corresponding antibodies. α-Tubulin immunoblotting here worked as the loading control. (b) Double knockout of *Ski* and *SnoN* rescues *in vitro Arkadia*-deficient Treg cell differentiation. T cells tranfected with the indicated gRNAs were cultured under Treg differentiation conditions for 3 days after 24h activation with plate-bound anti-CD3ε and anti-CD28 post electroporation. Shown is representative flow cytometry (left) and summary analysis of FOXP3 expression from technical replicates of one of 3 independent experiments. Statistical analyses were performed with unpaired t test. ns = not significant and *** *P* < 0.001. (c) Double knockout of *Ski* and *SnoN* rescues iTreg cell differentiation in *Arkadia*-deficient T cells *in vivo*. Upper panel, scheme of experimental design. Naïve T cells expressing Cas9, CD45.1/CD45.1 and *H. hepaticus* antigen-specific HH7-2 TCR were electroporated with gRNA vector for *Arkadia*, and control, *Arkadia*, or *Ski*-*SnoN*-*Arkadia*-triple targeting gRNAs were delivered into naïve T cells expressing Cas9, CD45.2/CD45.2 and HH7-2 TCR. Equal numbers of CD45.1/CD45.1 and CD45.2/CD45.2 electroporated cells were mixed and transferred into CD45.1/CD45.2 recipient mice colonized *H. hepaticus* (designated as groups 1, 2 and 3). To exclude microbiome variation among cages, group 3 mice were co-housed with mice from groups 1 and 2. To compare experimental groups, the ratios between co-transferred cells in each animal were calculated individually and combined for analysis with unpaired t test, ns = not significant, ** *P* < 0.01 and *** *P* < 0.001.

Because SKI and SnoN proteins are transcriptional repressors, we hypothesized that their inactivation may rescue the impairment of Treg cell differentiation in Arkadia-null T cells. We therefore expressed *Arkadia*-targeting gRNA with gRNAs targeting either *Ski, SnoN* or both in naive T cells from Cas9-expressing mice, followed by differentiation under Treg and Th17 conditions. Notably, targeting of both *Ski* and *SnoN*, but not of either gene alone, rescued Treg cell differentiation in Arkadia-deficient T cells, suggesting that SKI and SnoN have redundant roles in regulating the expression of FOXP3 in response to TGF-β (Fig. 4b). Consistent with their principal function of recruiting histone deacetylases^46^, deficiency of SKI and SnoN resulted in increased acetylation of histone H3K27 in the TGF-β-responsive *CNS1* of *Foxp3* (Extended Data Fig. 7b), indicative of transcriptional activation. By contrast, lack of SKI and SnoN had no significant influence on Th17 cell differentiation (Fig. 4b).

To investigate whether double knockout of *Ski* and *SnoN* rescues iTreg cell differentiation in *Arkadia*-null T cells *in vivo*, we delivered *Arkadia* or *Ski*-*SnoN*-*Arkadia*-triple targeting gRNAs into isotype-marked naïve T cells expressing Cas9 and the HH7-2 TCR that specifically recognizes *H. hepaticus* antigen^44^ (Fig. 4c). The isotype-distinct cells were mixed at a 1:1 ratio and transferred into mice colonized with *H. hepaticus*, followed by analysis of differentiated donor-derived cells in the LILP. Consistent with the *in vitro* differentiation results, deficiency of both SKI and SnoN rescued antigen-specific iTreg cell differentiation in Arkadia-deficient T cells. Taken together, our data suggest that TGF-β-activated Arkadia degrades SKI and SnoN, allowing for unopposed acetylation of histone H3 in the *Foxp3* locus and subsequent transcriptional activation.

## Discussion

The mammalian TGF-β superfamily is comprised of more than 30 members, including TGF-βs, activins, NODAL and inhibins, which transduce signals by way SMAD2/3, and bone morphogenetic proteins (BMPs), which signal through SMAD1/5/8^27^. TGF-β signaling has pleiotropic and indispensable roles in diverse biological processes, including embryonic development, immune responses, stem cell pluripotency and cell differentiation. Intriguingly, TGF-β signaling elicits seemingly conflicting responses in different cells and tissue settings. For example, TGF-β potently inhibits the growth of most cell types, including epithelial and endothelial cells, while stimulating growth in fibroblasts and smooth muscle cells. Additionally, in cancer, TGF-β can suppress pre-malignant tumorigenesis, yet promote metastasis^27^. This double-edged behavior of TGF-β signaling is present in many processes. The receptor-regulated SMAD effectors, SMAD2 and SMAD3, are phosphorylated by TGF-β receptor 1 and form hetero- or homo-trimers with SMAD4 to regulate the transcription of downstream genes. While SMAD2 and SMAD3 have some functional redundancy, they also have been shown to play distinct signaling roles. Similar to *Smad4*^-/-^ mice, *Smad2*^-/-^ mice die in utero, yet Smad3^-/-^ mice are viable and fertile but smaller in size. Further downstream in the signaling pathway, SKI and SnoN proteins act as functionally redundant suppressors of the TGF-β/NODAL pathways and are degraded by Arkadia. Interestingly however, only *Ski*-null mice are embryonic lethal while *SnoN* knockout mice have no described phenotype except for a mild defect in T-cell activation.

These seemingly inconsistent functions of TGF-β signaling components suggest still undiscovered intricacies to the fundamental signaling pathway. Three types of contextual determinants are considered to shape the responses of cells to TGF-β stimuli, including 1) the activation of canonical and non-canonical pathways by ligand-receptor pairing, 2) the factors cooperating with SMADs to regulate downstream transcription and 3) the epigenetic landscapes in cells^27^. Therefore, deciphering both the direct interactions and indirect regulatory mechanisms are important to fully understand the complexity of this pathway.

The roles of TGF-β in immune responses are also incompletely understood. Components of the canonical TGF-β pathway have essential roles in iTreg cell differentiation. These include TGF-β ligands, TGF-β receptors and SMAD4, but SMAD2 and SMAD3 are functionally redundant *in vivo^29^*. In this paper, we demonstrate that Arkadia is indispensable for Treg cell differentiation *in vitro*, yet its loss *in vivo* results in a relatively mild phenotype compared to the lack of TGF-β receptors. Mice with Arkadia deficiency in T cells did not develop autoimmune disease or severe spontaneous inflammation. However, they were more susceptible to pro-inflammatory conditions, which correlated with the reduction in iTreg cells. Arkadia ubiquitinates and promotes the degradation of transcriptional repressors, SKI and SnoN, to induce the expression of *Foxp3*. The reduction but not complete ablation of iTreg cells *in vivo* could be attributed to at least two possibilities. In addition to Arkadia, SKI and SnoN were also reported to be degraded by Anaphase-Promoting Complex (APC)^47, 48^ and Smurf2^49^. Although Arkadia-null cells failed to degrade SKI and SnoN with the treatment of TGF-β, we cannot exclude the possibility that APC and Smurf2 or other E3 ligases might contribute to degradation of SKI and SnoN *in vivo*. The other reason could be complex regulation of FOXP3 expression *in vivo*. SMAD3 and NFAT bind to the *CNS1* enhancer region, which is important for iTreg induction. The *CNS2* enhancer contains CREB and STAT5 binding sites, and is critical for maintenance of mature iTreg cells. Although SKI and SnoN bind with SMADs to inhibit FOXP3 transcription, the suppressive effect could be compensated by the presence of other transcriptional activators that are in part regulated by Arkadia-independent TGF-β receptor signaling

In contrast to Treg cells, while TGF-β and its receptors are vital for Th17 responses, SMADs play divergent and context-specific roles to downstream cell signaling. Lack of SMAD4 partially restored the expression of RORγt but not IL-17A in the absence of TGF-β, indicating both transcriptional suppression and activation-mediated by SMAD4. Moreover, mice lacking SMAD4 and TGFβR2 in T cells developed Th17-dependent EAE and expressed comparable IL-17A in spinal cord-infiltrating T cells^30^. Yet, in the LILP, we observed significantly reduced RORγt^+^FOXP3^-^ Th17 cells, suggesting different regulatory mechanisms in various tissues and environments. Dual deficiency of SMAD2/SMAD3 in T cells resulted in the loss of IL-17A production but comparable RORγt in response to IL6 plus TGF-β. The impaired expression of IL-17A induced by ablation of SMAD2/SMAD3 was restored in the presence of anti-IL-2 antibody, presumably through the loss of suppression in the *Il-17a* locus by STAT5^50^. Given the dispensable roles of SMAD2/SMAD3 in the induction of RORγt by TGF-β, an interesting question is whether SMAD2/SMAD3 also inhibit RORγt expression in the absence of TGF-β signaling, since SKI and SnoN also bind to SMAD2/SMAD3. This might potentially explain the partial restoration of RORγt expression in SMAD4-deficient T cells without TGF-β signaling. Alternatively, there may be non-canonical TGF-β signaling that regulates the expression of RORγt.

The lack of phenotype of Arkadia deficiency in Th17 cells raises some additional questions. We detected no obvious degradation of SKI and SnoN but also no defect of Th17 polarization in the Arkadia-deficient Th17 cells, which argues against SKI and SnoN negative regulation of TGF-β signaling through binding to SMADs. One possibility for differences seen in Th17 and iTreg differentiation might be the presence of distinct SMAD-containing complexes in the two programs. Alternatively, other unidentified, suppressors that are not substrates of Arkadia may negatively regulate RORγt expression. We have tested several suppressors involved in the TGF-β pathway, including *Smad7, Sip1, Snip1, Tgif1* and *Sin3a*, for their role in Th17 cell polarization. However, none of these showed an important role in the regulation of RORγt expression (data not shown). Therefore, the mechanism involved in TGF-β-mediated Th17 differentiation, which we find to be Arkadia-independent, remains incompletely understood.

To date, SKI and SnoN are the only widely accepted substrates of Arkadia. SnoN is efficiently degraded when it forms a complex with both Arkadia and phosphorylated SMAD2/SMAD3^31^. However, little is known about the regulation of Arkadia and its roles in different cell types that respond to Activin, NODAL and TGF-β stimuli. Furthermore, we cannot rule out the possibility that other proteins involved in TGF-β signaling are degraded by Arkadia. Given that the upstream signals are conserved in many cell types, further dissection of the downstream TGF-β signaling components and their relative roles in different biological processes will be of utmost importance for future studies. Specifically, this work highlights that modulation of Arkadia might provide a platform to gain unique insight into the regulation of various physiological processes by TGF-β superfamily members.

## Acknowledgements

We acknowledge support from S.Y. Kim and the NYU Rodent Genetic Engineering Laboratory (RGEL) for generating *Arkadia* conditional knockout mice, the Experimental Pathology Research Laboratory of NYU Medical Center for Histologic processing and imaging. Both laboratories were partially supported by the Cancer Center Support Grant P30CA016087 at NYU Langone’s Laura and Isaac Perlmutter Cancer Center. This work was supported by the Irvington Institute fellowship program of the Cancer Research Institute (H.X.); the Helen and Martin Kimmel Center for Biology and Medicine (D.R.L.); the Howard Hughes Medical Institute (D.R.L.); and National Institutes of Health grant R01AI080885 (D.R.L.).

## Author contribution

H.X. and D.R.L. designed experiments. H.X. performed experiments and analyzed the data. L.W. developed the method to express gRNA in naïve T cells by electroporation. H.N. performed blinded histology scoring on colon sections. V.R. helped with EAE and oral gavage of bacteria. V.E. provided advice on Arkadia experiments and helped validate experimental reagents. H.X., K.M. and D.R.L. wrote the manuscript with input from the co-authors. D.R.L. supervised the research.

## Methods

### Mice

Mice were bred and maintained in the animal facility of the Skirball Institute (New York University School of Medicine) under specific pathogen-free conditions. All animal procedures were performed in accordance with guidelines by the Institutional Animal Care and Usage Committee of New York University School of Medicine or National Institute of Allergy and Infectious Diseases (NIAID). C57BL/6, *Cd4^Cre^* (*Tg*(*Cd4-cre*)*1Cwi/BfluJ*), *Cd45.1* (*B6.SJL-Ptprc^a^ Pepc^b^/BoyJ*) and Rosa26-Cas9 knockin (*Gt(ROSA)26Sor^tm1.1(CAG-cas9*,-EGFP)Fezh^*/J) mice were obtained from Jackson Laboratories or Taconic Farm. *Smad4^fl/fl^* (*Smad4^tm2.1Cxd^*/J) and *Tgfbr2 ^fl/fl^* (*Tgfbr2^tm1Karl^*/J) mice were purchased from Jackson Laboratories and bred with *Cd4^Cre^* mice to give the *Smad4* and *Tgfbr2* double conditional knockout mice. HH7-2 TCR transgenic mice were previously described^44^. *Arkadia^fl/fl^* were generated by insertion of two LoxP sites flanking Exon13, and were further bred with *Cd4^Cre^* mice for conditional knockout of *Arkadia* in T cells. All experiments were performed with genetically modified mice and control littermates with matched sex and age. If not specified, mice used in all experiments were 6 to 12 weeks old when treated with different stimuli.

### Cell lines, antibodies for flow cytometry and western blotting

The following monoclonal antibodies (clone ID) were used for flow cytometry (Extended Data Table III): CD4 (RM4-5), CD8 (53-6.7), CD25 (PC61.5), CD44 (IM7), CD45.1 (A20), CD45.2 (104), CD62L (MEL-14), TCRβ (H57-597), TCR Vβ6 (RR4-7), FOXP3 (FJK-16s), GATA3 (TWAJ), Helios (22F6), RORγt (B2D), T-bet (eBio4B10), IL-17A (eBio17B7) and IFNγ (XMG1.2). 4′,6-diamidino-2-phenylindole (DAPI) or Live/ dead fixable blue (ThermoFisher) was used to discriminate live/dead cells. Anti-acetyl-H3K27 (D5E4, Cell Signaling Technology) and IgG isotype control (DA1E, Cell Signaling Technology) antibodies were employed in ChIP assay. The following antibodies were used for western blotting: SKI (Santa Cruz, sc-33693), SnoN (Santa Cruz, sc-9141), α-tubulin (Santa Cruz, sc-69970). Protein concentration of cell lysate was determined by Bradford and Lowry Methods (Bio-Rad) and 70 µg proteins were loaded for western blotting. *SnoN^-/-^* and *Ski^-/-^* MEF are described in a preprint on biorxiv (doi: https://doi.org/10.1101/487371). MEF were cultured in DMEM medium containing 10% FBS, 20 mM L-glutamine and 1× Penicillin/Steptomycin.

### *In vitro* T cell differentiation

Naïve CD4^+^ T cells were defined as CD4^+^ CD8^-^ CD25^-^ CD44^-^ CD62L^+^ cells and isolated on an Aria II (BD Biosciences) cell sorter. For polarization of T cells *in vitro*, goat-anti-hamster IgG (MP Biomedicals) was diluted (1:20) in PBS to coat 96-well plates overnight. T cells were activated by adding hamster anti-CD3ε (0.25 µg/mL) and anti-CD28 (1 µg/mL) monoclonal antibodies. Polarizations were performed in 300 µL RPMI medium supplemented with exogenous cytokines for different cell lineages: Th0, 100 unit/mL hIL-2 (Peprotech) and 10 µM TGFβR1 inhibitor SB525334 (selleckchem); Th1, 100 unit/mL hIL-2, 10 ng/mL IL-12 (Peprotech) plus 2 µg/mL anti-IL-4 antibody (BioXcell, Clone: 11B11); Th17, 0.2 ng/mL TGF-β (Peprotech), 20 ng/mL IL-6 (Ebioscience), 2 µg/mL anti-IL-4 and anti-IFNγ antibody (BioXcell, Clone: XMG 1.2). 10 µM SB525334 was used to replace TGF-β to inhibit TGF-β signaling if required; iTreg, 500 unit/mL hIL-2 and 5 ng/mL TGF-β. For Th0 and Th1 cell polarization, 2 × 10^4^ cells were plated in each well and transferred to un-coated 96-well plates after two days, followed by another 1 or 2 days of polarization. For Th17 and iTreg cell polarization, 1 × 10^4^ cells were plated in each well and polarized for 3 days. For re-stimulation and cytokine analysis, cells were incubated at 37 °C for 3h in fresh medium with 50 ng/mL phorbol 12-myristate 13-acetate (PMA, Sigma), 500 ng/mL ionomycin (Sigma) and 1 × GolgiStop (BD).

For transcription factor staining, cells were stained for surface markers prior to nuclear factor staining following the manufacturer’s protocol (00-5523, eBioscience). For intracellular cytokines staining, Cytofix/Cytoperm buffer set (BD Biosciences) was used. Analysis of stained cells was performed on a LSR II (BD Biosciences) flow cytometer, and data were analyzed on FlowJo platform (BD).

### Package of retrovirus and viral infection of T cells

The retroviral plasmid, pSIN-U6-sgRNA-EF1a-Thy1.1-P2A-Neo or MSCV-IRES-Thy1.1, was used for viral packaging and expression of gRNA or Arkadia, respectively. Plat-E cells (Cell Biolabs) and jetPRIME transfection reagent (Polyplus) were used to produce retrovirus, which was harvested at 48h after transfection. Before viral transduction in 96-well plates, 1 × 10^4^ naïve T cells in 100 µL medium were activated by anti-CD3ε and anti-CD28 cross-linking for 24 hours. 100 µL retrovirus-containing medium was added to each well for viral transduction in the presence of 5 µg/mL polybrene. Cells were centrifuged at 2200 rpm for 90 minutes at room temperature, followed by culture for 12h in a cell incubator. Cells were then polarized under required conditions for 3 days.

### Isolation of lymphocytes from SILP or LILP

Intestines were collected and carefully inspected to remove Peyer’s patches in small intestines and caecal patches in large intestines. After cleaning, intestines were sequentially treated with Hank’s Balanced Salt Solution (HBSS) containing 1 mM DTT once at room temperature for 15 minutes, and HBSS containing 5 mM EDTA twice at 37 °C for 30 minutes total time to remove epithelial cells. Tissues were minced in RPMI complete medium containing collagenase (1 mg/mL, Roche), DNase I (100 μg/mL, Sigma), dispase (0.05 U/mL, Worthington), followed by shaking for 50 minutes at 37 °C. Leukocytes were isolated by 40%/80% Percoll gradient centrifugation (GE Healthcare). For re-stimulation *ex vivo*, isolated leukocytes were suspended in fresh medium containing 50 ng/mL PMA, 500 ng/mL ionomycin and 1 × GolgiStop and incubated at 37 °C for 3h before staining. Thymus, lymph node and spleen were mechanically disrupted directly on a 70 μm filter, followed by removal of blood cells with ACK lysis buffer.

### *H. hepaticus* culture and oral infection with SFB and *H. hepaticus*

*H. hepaticus* was inoculated on blood agar plates (TSA with 5% sheep blood, Thermo Fisher) and cultured in a hypoxia chamber (Billups-Rothenberg) at 37 °C for 4 to 5 days. The anaerobic atmosphere was achieved by aeration of gas mixture composed of 80% nitrogen, 10% hydrogen, and 10% carbon dioxide (Airgas). Before animal inoculation, *H. hepaticus* were suspended in Remel Brucella Broth (Thermo Fisher). The concentration of bacteria was adjusted to an optical density of 1 to 1.5 measured at a wavelength of 600 nm. Each mouse was administered 0.2 mL/dose of bacterial suspension on day 0 and day 4 by oral gavage. For SFB infection, one fecal pellet from Taconic mice was suspended in 200 µL PBS and passed through a 100 µm filter. The filtrate was given to one mouse by oral gavage. Each mouse received two doses totally with an interval of 3 days. T lymphocytes of intestinal lamina propria were recovered and analyzed 3 to 4 weeks after the last gavage.

### Bone marrow transfer

Recipient mice (CD45.1/CD45.2) were irradiated with 500 cGy + 500 cGy with an interval of 5 hours one day before bone marrow transfer (Day 0). On day 1, bone marrows were collected from congenic mice (CD45.1/CD45.1), control or *Arkadia* conditional knockout mice (CD45.2/CD45.2). Erythrocytes in bone marrow were removed by ACK lysis buffer and T cells were depleted using anti-CD90.2 magnetic beads (Miltenyi), followed by determination of cell number. Equal numbers of bone marrow cells from control or *Arkadia* conditional knockout mice (CD45.2/CD45.2) were mixed with cells from congenic mice (CD45.1/CD45.1), respectively, followed by transfer of 3 × 10^6^ mixed cells into each lethally irradiated CD45.1/CD45.2 recipient mouse through retroorbital injection. Antibiotics (1 mg/mL sulfamethoxazole and 0.2 mg/ mL trimethoprim) were added to the drinking water and maintained for 2 weeks. CD4^+^ TCRβ^+^ T lymphocytes from LILP were stained for RORγt and FOXP3 3 months after transfer.

### IL-10RA blockade and histology scoring on colon sections

Under SPF conditions, *Arkadia* conditional knockout mice and control littermates were cohoused after birth and intraperitoneally administered one dose of 1 mg/mL anti-IL-10RA antibody at 10 to 12 weeks old. About 5 weeks later, mice were sacrificed to isolate lamina propria lymphocytes or collect colonic samples for histology analysis. With *H. hepaticus*-colonized mice, one dose of 0.25 mg/mL anti-IL-10RA antibody was administered intraperitoneally. Mice were sacrificed to isolate LILP lymphocytes or collect colons for histology analysis 3 weeks after the IL-10RA blockade. Colonic samples were sequentially fixed in 4% paraformaldehyde (Electron Microscopy Science, Hatfield USA), embedded in paraffin and sectioned. Colon sections were stained with hematoxylin and eosin and scored for histopathology in a blinded fashion. Total score of colonic inflammation comprised individual scores from 4 categories: 1) Number of goblet cells per High Power Field (HPF) (Score of 1: 11 to 28 goblet cells per HPF; Score of 2: 1 to 10 goblet cells per HPF; Score of 3: less than 1 goblet cell per HPF). 2) Submucosa edema (Score of 0: No edema; Score of 1: Mild edema accounting for < 50% of the diameter of the entire intestinal wall; Score of 2: moderate edema involving 50%–80% of the diameter of the entire intestinal wall; Score of 3: profound edema involving > 80% of the diameter of the entire intestinal wall). 3) Depth of lymphocytes infiltration (Score of 0: No infiltrate; Score of 1: Infiltration above muscularis mucosae; Score of 2: Infiltration extending to- and including submucosa; Score of 3: Transmural Infiltration). 4) Epithelial changes (Score of 0: No changes; Score of 1: Upper third or only surface epithelial missing; Score of 2: Moderate epithelial damage with intact base of crypts; Score of 3: Loss of crypts).

### Experimental allergic encephalomyelitis (EAE)

On day 0, the following reagents were prepared for injection: pertussis toxin (200 ng/mouse in 200 μL PBS), Mycobacteria tuberculosis H37RA suspended in CFA with a final concentration of 2 mg/mL (100 μL for each mouse) and MOG peptide (200 μg/mouse in 100 μL PBS). Equal volume of Mycobacteria tuberculosis H37RA-CFA was mixed with MOG/PBS in a 5 mL protein Lobind tube (Eppendorf), followed by emulsification in ice water through sonication. The mixture was centrifuged at 3000 rpm for 2 minutes to confirm emulsion (no phase separation after centrifugation). 200 μL mixture was injected into each mouse subcutaneously with 27 G½ needle (BD). On day 2, mice were administered a second dose of pertussis toxin (200 ng/mouse in 200 μL PBS). The disease score was measured daily one week after the last pertussis toxin injection. A scoring system from 0 to 10 was applied to measure the severity of disease: 0, no clinical signs; 1, limp tail; 2, paralyzed tail; 3, hind limb paresis; 4, one hind limb paralysis; 5, both hind limbs paralysis; 6, fore limb weakness; 7, one fore limb paralysis; 8, both fore limbs paralysis; 9, moribund; 10, death. The clinical scores were further analyzed by two-way ANOVA to determine difference between two mouse lines.

### Knockout of genes by electroporation

All gRNAs were expressed by transduction of retrovirus packaged with pSIN-U6-sgRNA-EF1a-Thy1.1-P2A-Neo plasmids. The gene-editing efficiencies of gRNAs were first confirmed by western blotting or staining of targeted proteins in polarized T cells. For knockout of genes by electroporation (4D-Nucleofector, Lonza), PCR products were amplified with primers (Extended Data Table IV) flanking the hU6 promoter, sgRNA and sgRNA scaffold in pSIN-U6-sgRNA-EF1a-Thy1.1-P2A-Neo plasmids. For each gRNA-expressing DNA, 400 µL PCR products were purified by Promega Wizard Gel/PCR purification columns, followed by NaAc-Ethanol purification. Briefly, DNA products eluted from columns were mixed with 1/10 volume of 3 M NaAc pH5.2 and 2.5-fold volume of 100% ethanol. The mixture was placed at -20 °C overnight and centrifuged for 30 minutes to precipitate DNA. The supernatant was removed and DNA was washed twice with 70% ethanol (pre-cold at -20 °C). 30 µL water was added to dissolve air-dried DNA. DNA concentration was determined by NanoDrop (the concentration was typically around 0.5 µg/µL). Naïve T cells expressing Cas9 were washed once with 1 mL pre-warmed recovery medium (Mouse T Cell Nucleofector Medium, Lonza) before electroporation. 2 × 10^6^ cells were suspended in 100 µL buffer (P3 Primary Cell 4D-Nucleofector X Kit, Lonza), mixed with 4 µg DNA and transfected using the 4D-Nucleofector program DN-100. After electroporation, cells were suspended in 500 µL recovery medium and incubated in the incubator for 30 minutes. 2 × 10^4^ cells were then seeded into each well of 96-well plates and incubated with plate-bound anti-CD3ε and anti-CD28 cross-linking for 24h to ensure gene editing, followed by T cell differentiation for 3 to 4 days.

### Chromatin immunoprecipitation (ChIP)

As described above, plasmids encoding control, *Arkadia, Ski* or *SnoN* targeting gRNAs were delivered into naïve T cells expressing Cas9 through electroporation, followed by 24h activation and polarization for 3 days *in vitro.* 4 µg DNA was used for each electroporation, in which 1.33 µg DNA was used for each gene when targeting three genes. Control gRNA was used to achieve a total of 4 µg DNA, when necessary, for example 2.67 µg control DNA plus 1.33 µg Arkadia-targeting DNA for Arkadia knockout.Polarized Treg cells were then harvested for ChIP analysis, in which anti-acetyl-H3K27 and isotype control antibodies were used to precipitate targeted DNA fragments (#9005, cell signaling Technology). Enrichment of genomic DNA fragments by ChIP was validated by Real-time PCR (LightCycler® 480, Roche) with primers (Extended Data Table IV) targeting the *Foxp3 CNS1* region. Primers (#7015, cell signaling Technology) targeting ribosomal protein L30 functioned as the control for the ChIP assay.

### Rescue of Arkadia deficiency *in vivo*

The protocol of electroporation to deliver DNA was the same as that for the ChIP assay. Briefly, plasmids encoding *Arkadia*-targeting gRNA were delivered into naïve T cells expressing Cas9, CD45.1/CD45.1 and HH7-2 TCR. Meanwhile, plasmids encoding control, *Arkadia* or *Ski*-*SnoN*-*Arkadia*-triple targeting gRNAs were delivered into naïve T cells expressing Cas9, CD45.2/CD45.2 and HH7-2 TCR. Equal numbers of CD45.1/CD45.1 and CD45.2/CD45.2 naïve cells (50,000 cells each) were mixed and designated as groups 1, 2 and 3 (Fig. 4c). To exclude microbiome variation among cages, group 3 was used as internal control to groups 1 and 2. The mixed cells were re-suspended in 100 µL medium containing 2% FBS and transferred by retroorbital injection into CD45.1/CD45.2 recipient mice colonized with *H. helicobacter*. Expression of RORγt and FOXP3 was examined in CD4^+^ TCRβ^+^ T lymphocytes from LILP 2 weeks after cell transfer.

### Statistical analysis

We used two-tailed *P*-value calculated by unpaired t test (Welch’s t test) to determine the differences between experimental groups. In experiments with co-transferred T cells or bone marrows, the ratio in each mouse between the co-transferred cells was calculated individually and combined together for unpaired t test. For EAE disease comparison, the clinical scores were analyzed by two-way ANOVA. *P*-value less than 0.05 was reported as significant difference, ns = not significant, * *P* < 0.05, ** *P* < 0.01, *** *P* < 0.001, and **** *P* < 0.0001. Detailed number of replicates and representative data can be found in the figure legends.

**Extended Data Figure 1.**
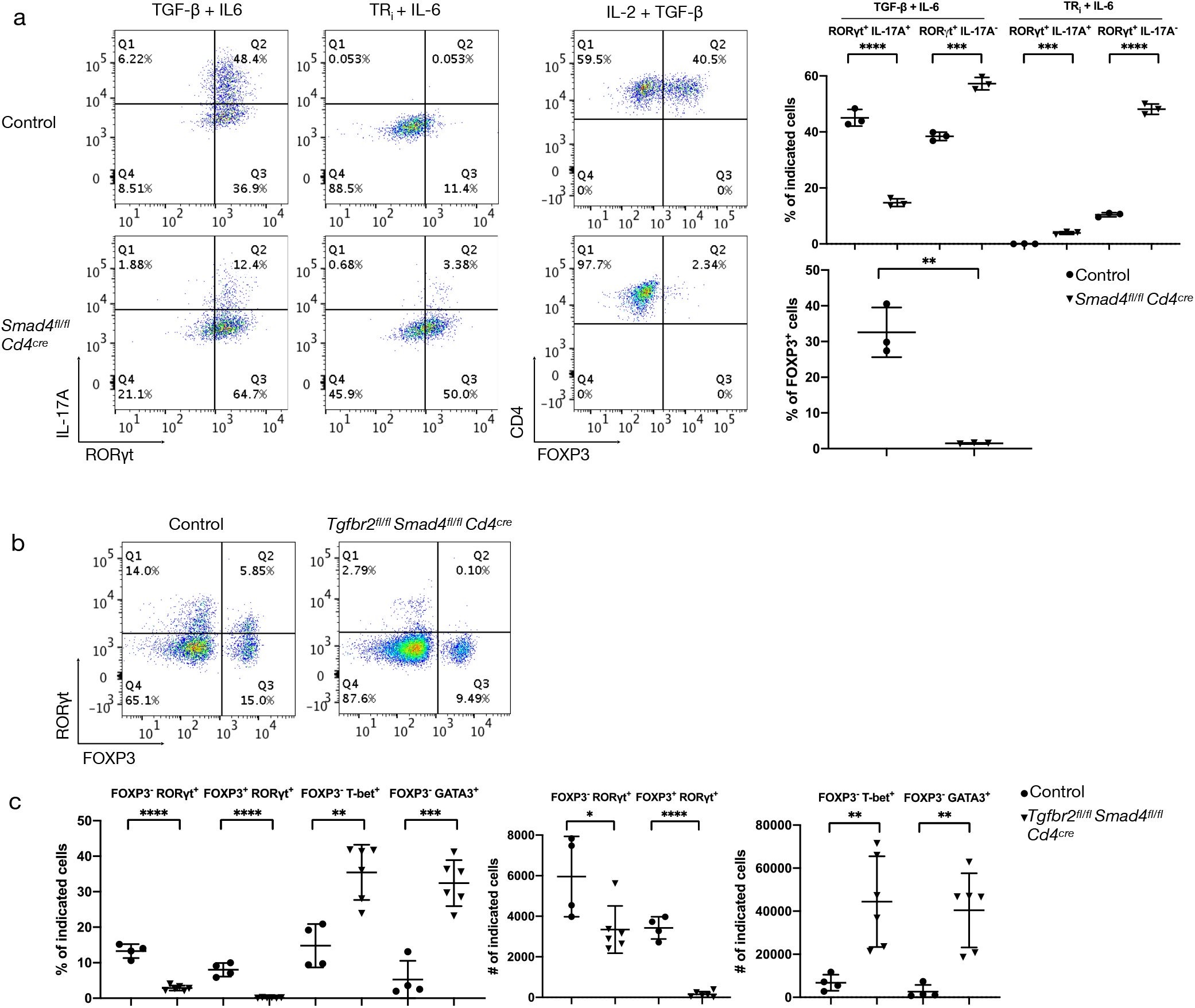
Differential requirement of SMAD4 in Treg and Th17 cell differentiation (a) Role of SMAD4 in Th17 and Treg cell differentiation. Representative flow cytometry panels of T cells from WT and conditional *Smad4* mutant mice, differentiated *in vitro* under Th17 and Treg conditions, or with blockade of TGF-β signaling with TR_i_ (SB525334), an inhibitor of TGFβR1, (left panels). Right panel, representative data of two independent experiments of T cells differentiated under the indicated conditions. *P*-values were calculated by unpaired t test. (b) Representative RORγt and FOXP3 staining of CD4^+^ TCRβ^+^ T lymphocytes from LILP of control littermates (with at least one allele expressing intact SMAD4 and TGFBRII) and *Tgfbr2^fl/fl^ Smad4^fl/fl^ Cd4^cre^* conditional double knockout mice (4 to 6 week old) kept under SPF conditions. (c) Proportion and total cell number of indicated T cell populations in the LILP from control and *Tgfbr2^fl/fl^ Smad4^fl/fl^ Cd4^cre^* mice, as shown in (b). Differences were determined by unpaired t test, * *P* < 0.05, ** *P* < 0.01, *** *P* < 0.001, and **** *P* < 0.0001.

**Extended Data Figure 2.**
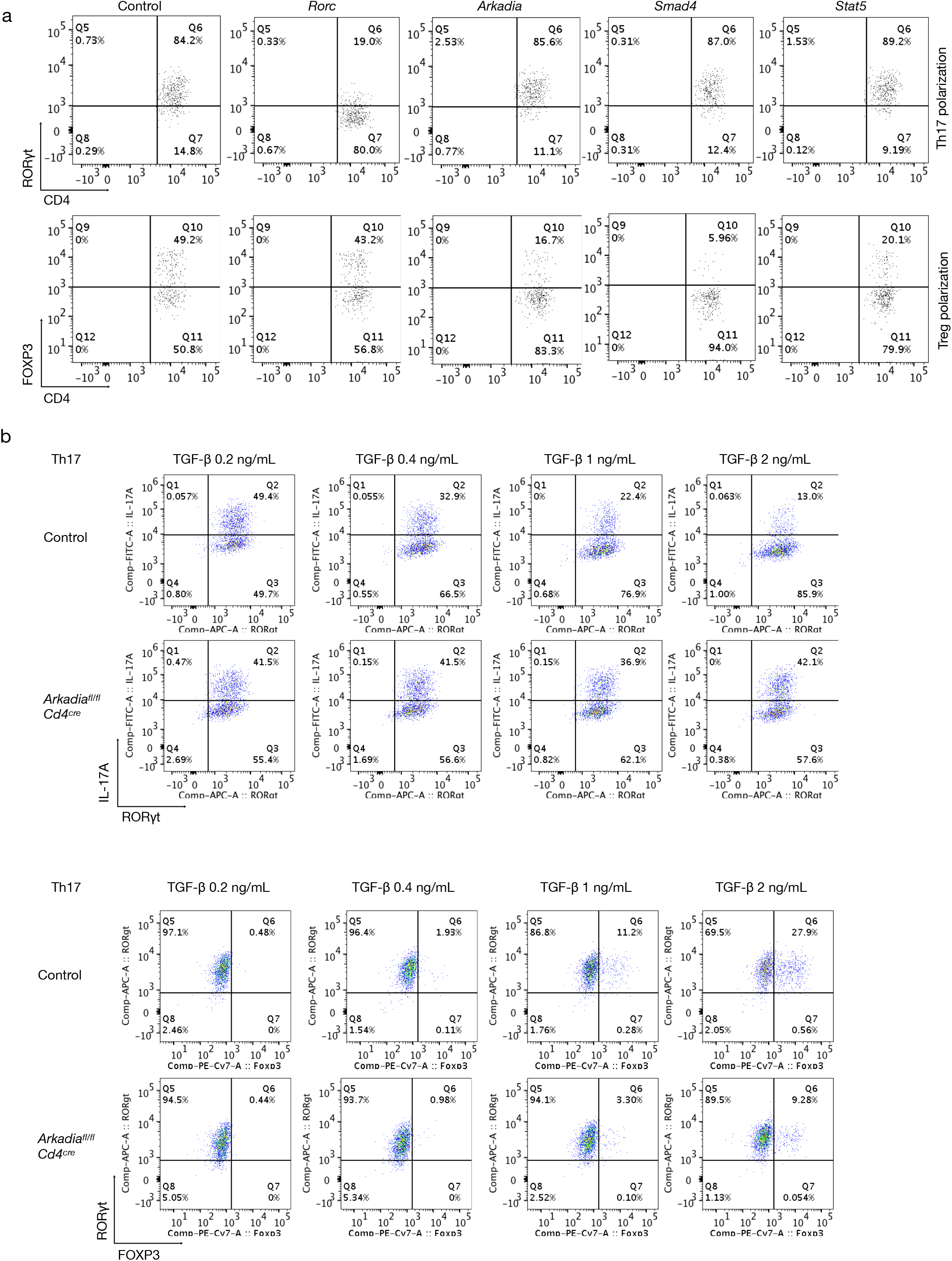
Arkadia requirement for in vitro differentiation of Treg, but not Th17, cells. (a) Flow cytometry analysis of CD4^+^ T cells differentiated *in vitro* under Th17 and Treg conditions following transduction of retroviruses expressing the indicated gRNAs. Naïve T cells were from Cas9 transgenic mice, and analysis was performed following gating on the Thy1.1 transduction marker. (b) Flow cytometry to assess IL-17A/ RORγt (top panels) and RORγt/FOXP3 (lower panels) expression in WT and *Arkadia^f/f^ Cd4^Cre^* T cells polarized in the presence of different concentration of TGF-β plus 20 ng/mL IL-6. IL-17A, RORγt and FOXP3 were stained as indicated.

**Extended Data Figure 3.**
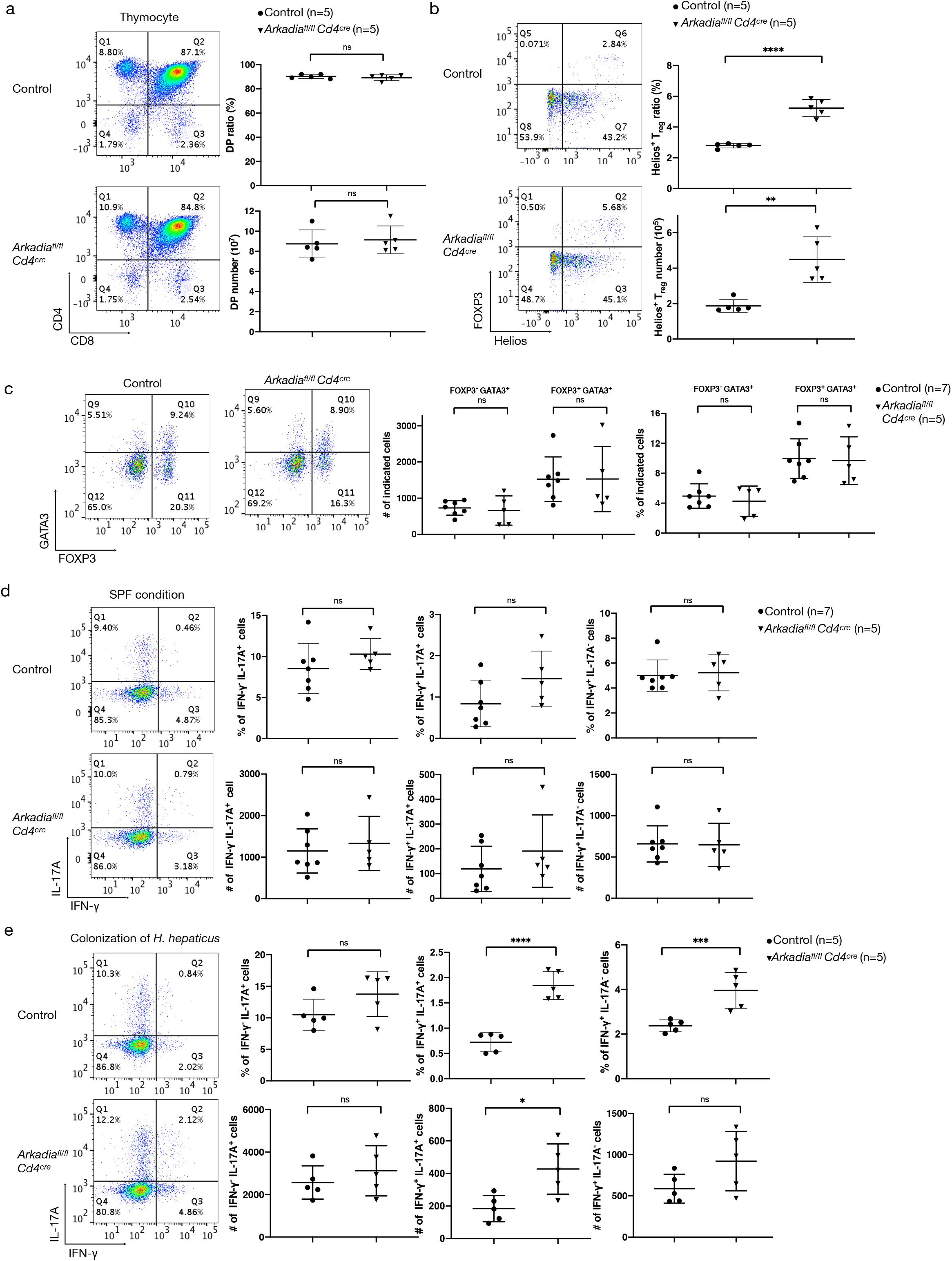
Loss of Arkadia in T cells results in elevated thymic Treg cells, but no perturbations in the intestinal T cell compartment of SPF mice or mice colonized with *H. hepaticus*. (a) Analysis of thymocyte subsets from control and *Arkadia* conditional knockout mice. Representative flow cytometry panels (left), proportions and total numbers of double positive thymocytes from multiple mice (right). (b) Representative flow cytometry (left) and nTreg proportions and numbers among gated CD4 single positive thymocytes of control and *Arkadia* conditional knockout mice. (c) Analysis of Treg cells among CD4^+^ TCRβ^+^ T lymphocytes from LILP of control and *Arkadia* conditional knockout mice under SPF conditions. Representative flow cytometry for GATA3 and FOXP3 (left) and cell proportions and numbers of indicated T lymphocyte subpopulations (right). (d, e) INF-γ and IL-17A production by CD4^+^ TCRβ^+^ T lymphocytes from LILP of control and *Arkadia* conditional knockout mice in SPF conditions (d) and after stable colonization with *H. hepaticus (e)*. Data for (c) and (d) are from 1 of 3 independent experiments, with n = 13 in the 3 experiments. Data for (e) are representative of 2 independent experiments (n = 10). (a to e) Unpaired t test was used for statistical analysis. ns = not significant, * *P* < 0.05, ** *P* < 0.01, *** *P* < 0.001, and **** *P* < 0.0001.

**Extended Data Figure 4.**
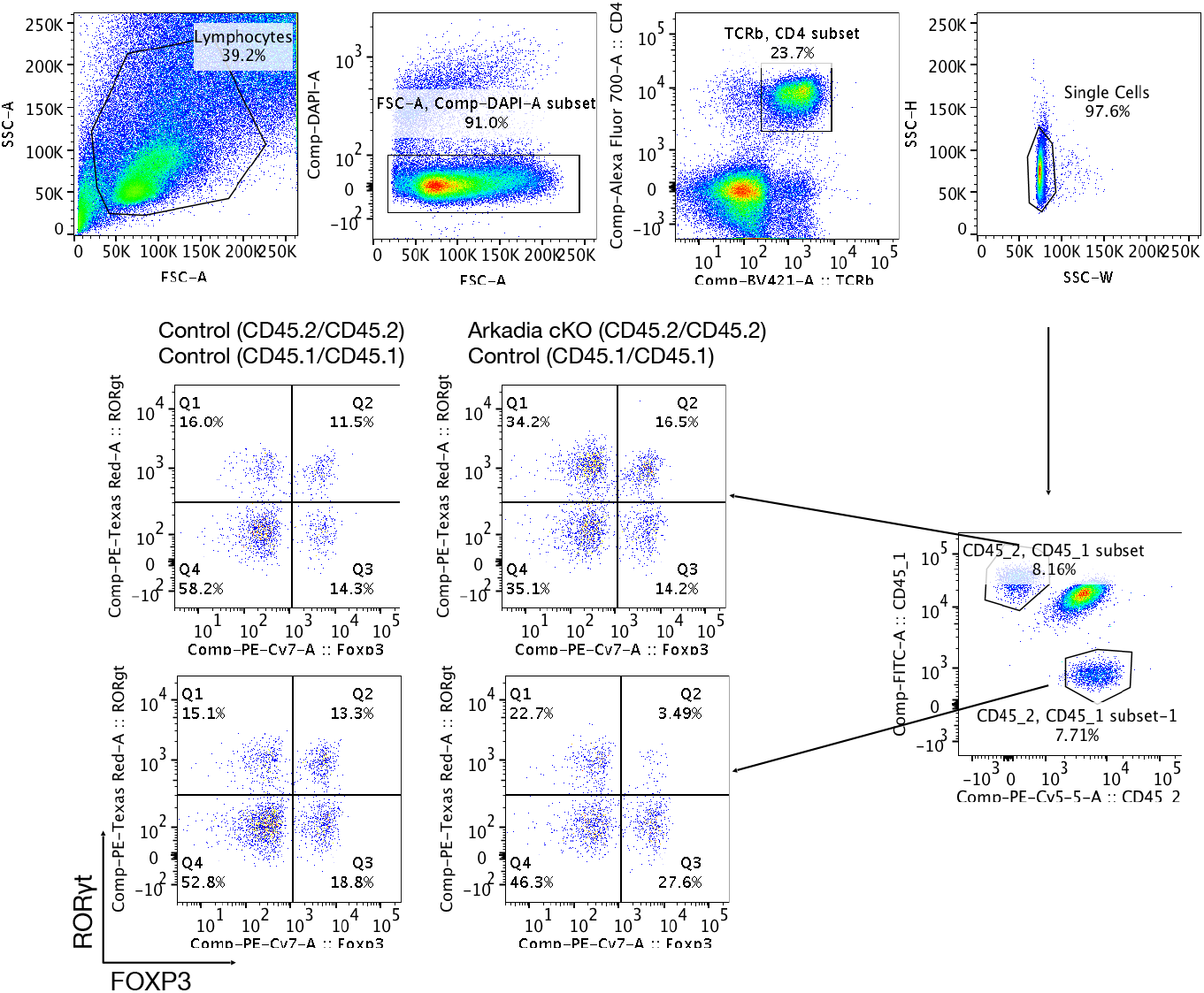
Gating strategy for the mixed bone marrow chimera experiments Gating to distinguish the donor-derived CD4^+^ TCRβ^+^ T lymphocytes from LILP of mice receiving equal numbers of bone marrow cells from either control or *Arkadia* conditional knockout mice (CD45.2/CD45.2) and CD45.1/CD45.1 mice (WT) mice (see Fig. 2c). Representative flow cytometry panels for control:WT and *Arkadia^f/f^ Cd4^Cre^*:WT donors are shown.

**Extended Data Figure 5.**
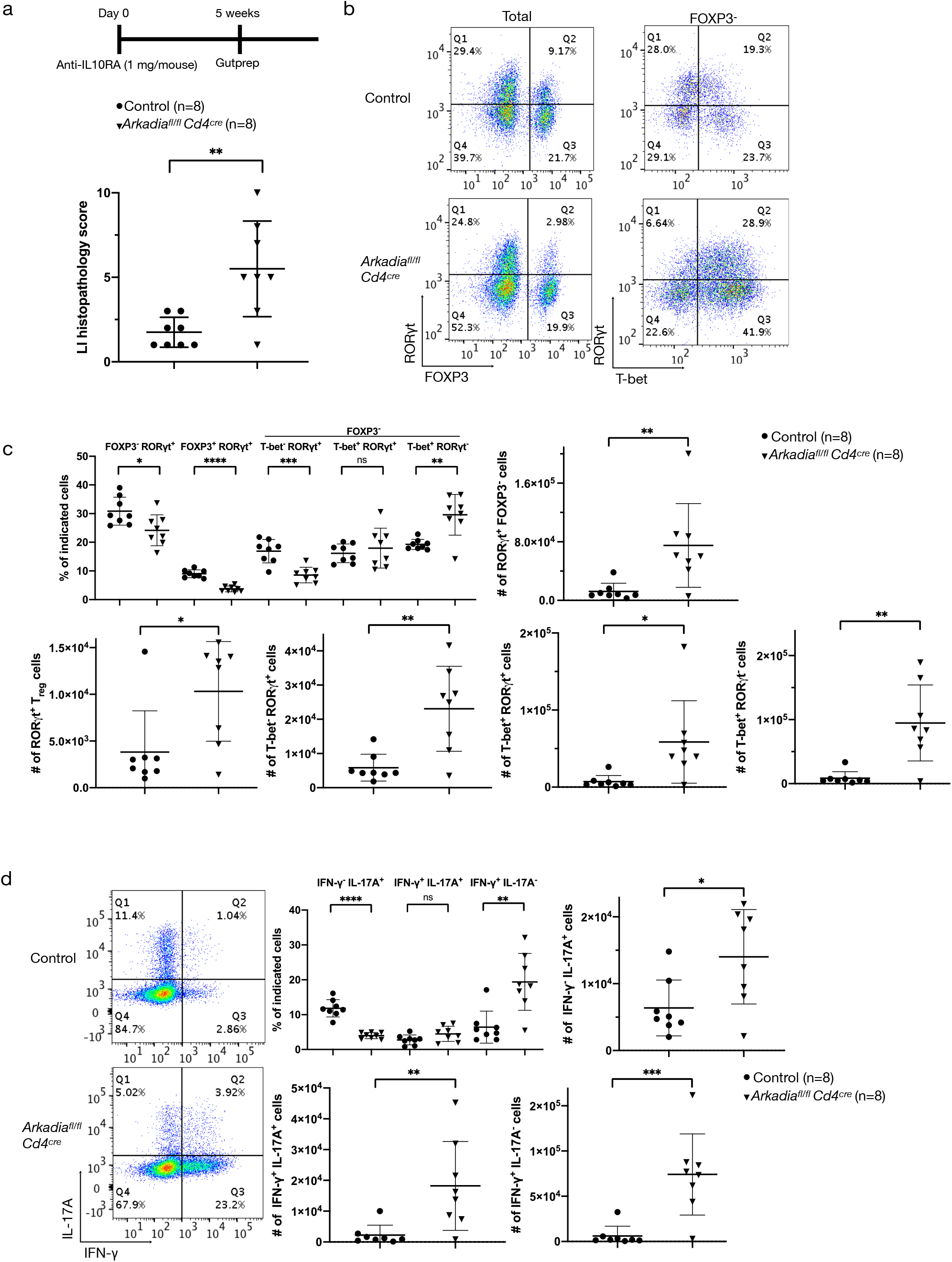
Arkadia-deficient mice are susceptible to immune system perturbation (a) Scheme of IL-10RA blockade under SPF conditions and analysis of colitis scores in colon of control and *Arkadia* conditional knockout littermates (n=8). (b-d) Representative flow cytometry and indicated T cell subpopulations (proportions and numbers) following IL-10RA blockade b, Representative RORγt, FOXP3 and T-bet expression in CD4^+^ TCRβ^+^ T lymphocytes isolated from LILP of indicated mice. c, proportions and numbers of indicated T lymphocyte subpopulations. d, representative flow cytometry analysis of INF-γ and IL-17A expression (left) and frequencies of cells in each biological replicate (right). Unpaired t test was used for statistical analysis. ns = not significant, * *P* < 0.05, ** *P* < 0.01, *** *P* < 0.001, and **** *P* < 0.0001.

**Extended Data Figure 6.**
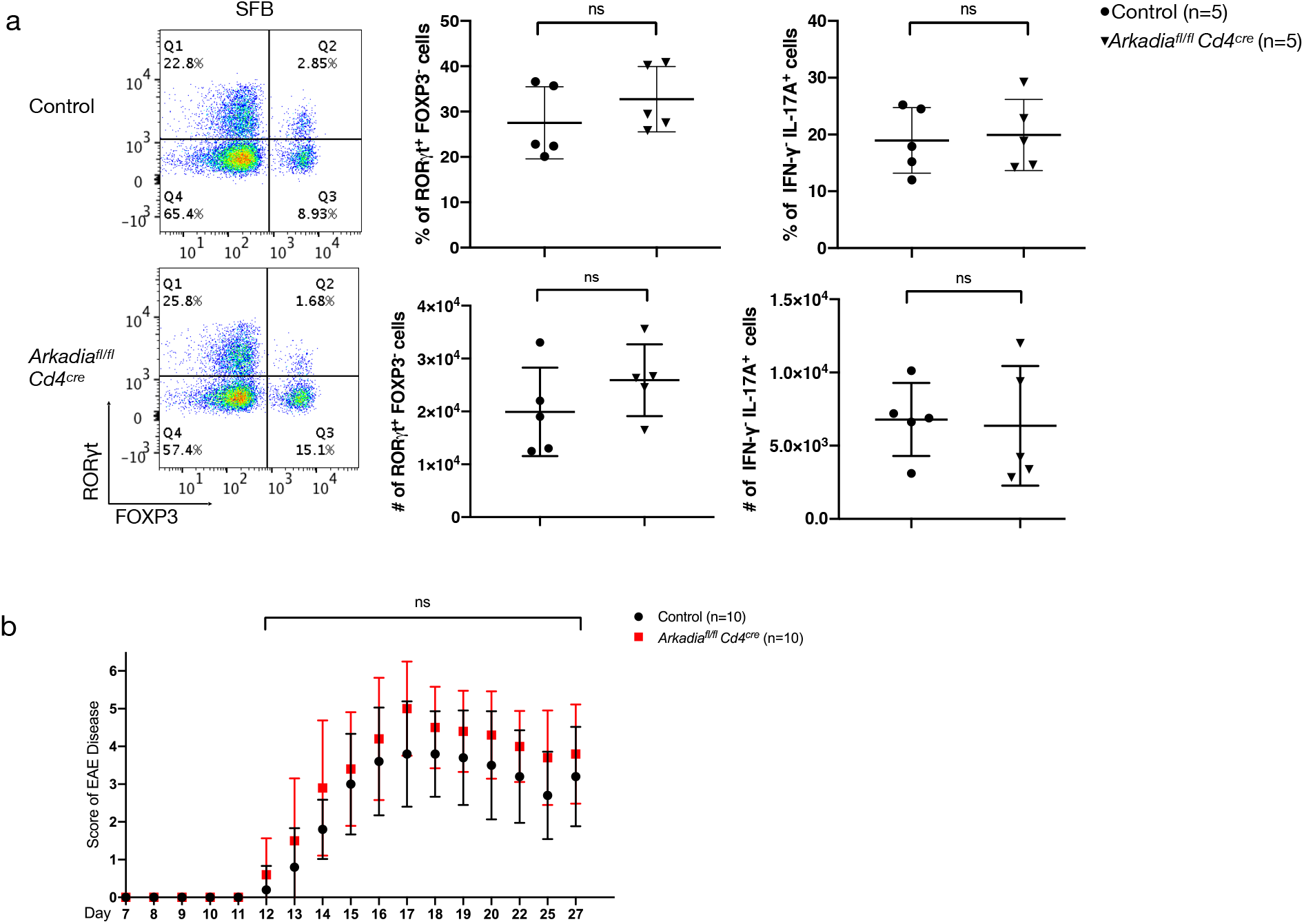
Arkadia does not influence SFB-dependent Th17 cell differentiation or Th17 cell-mediated EAE (a) Representative flow cytometry panel and analysis of RORγt, FOXP3 and effector cytokine expression in T lymphocytes from small intestine lamina propria (SILP) of mice colonized with segmented filamentous bacteria (SFB). (b) EAE disease score in control and *Arkadia* conditional knockout mice. The EAE clinical scores were analyzed by two-way ANOVA. ns = not significant.

**Extended Data Figure 7.**
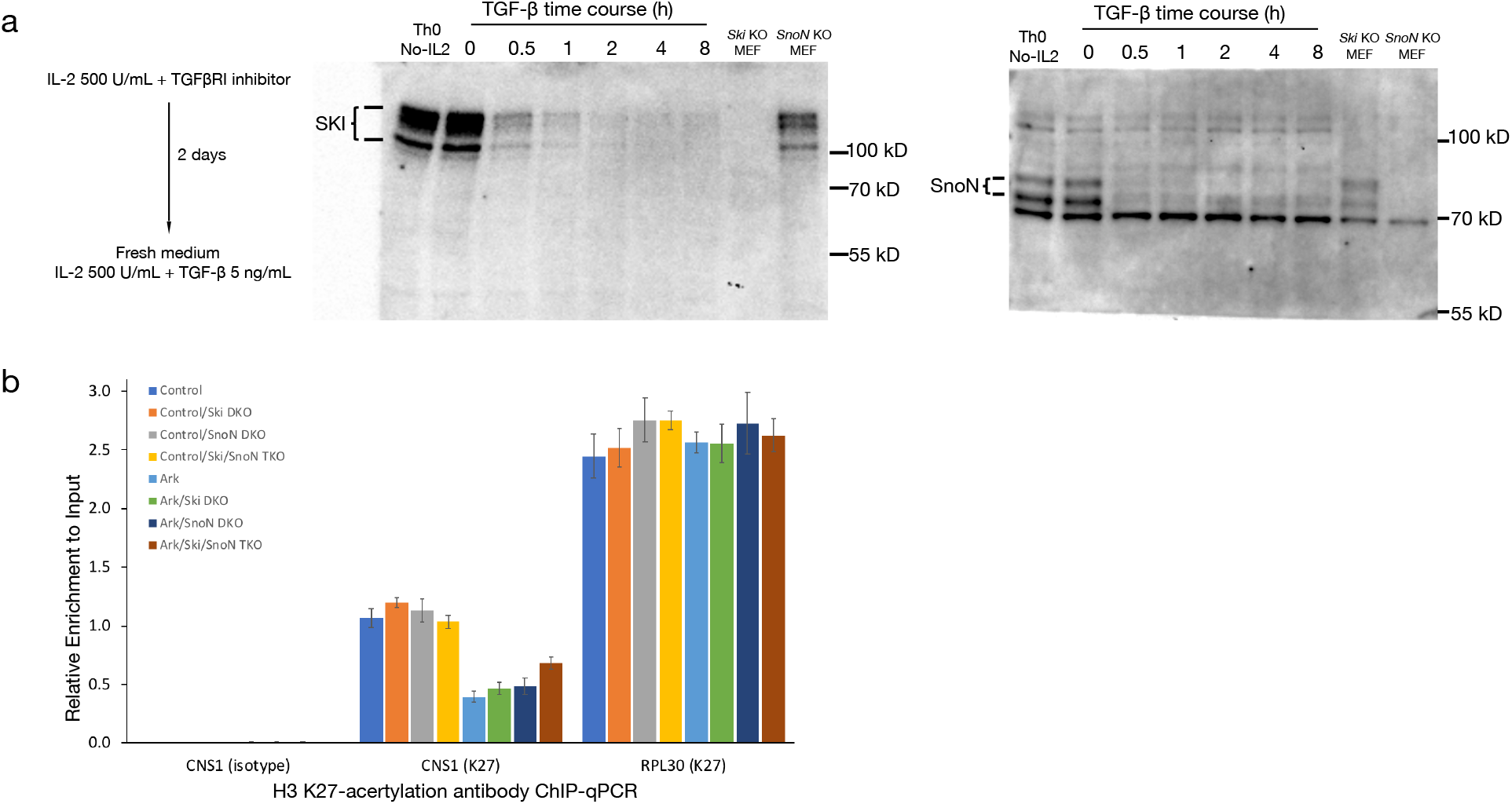
SKI and SnoN play redundant roles in the Arkadia-mediated signal cascade. (a) Validation of SKI and SnoN antibodies. T cells were polarized by plate-bound anti-CD3ε and anti-CD28 for 2 days in the presence of TGFβRI inhibitor and IL-2. T cells were then treated with fresh medium containing both IL-2 and TGF-β for indicated times before collection for western blotting. Cell lysates from SKI or SnoN-deficient MEFs were used as control. (b) The Arkadia-SKI/SnoN axis regulates FOXP3 expression through histone modification at the *Foxp3* locus. Naïve T cells expressing Cas9 were transduced with vector expressing sgRNAs to target *Arkadia, Ski* or *SnoN* (and with a non-targeting control), and after 24h activation they were cultured in differentiation conditions for 3 days. Polarized Treg cells were then harvested for ChIP analysis using anti-acetyl-H3K27 or isotype control antibodies. Enrichment of genomic DNA fragments by ChIP-qPCR was validated by primers targeting the *Foxp3 CNS1* region. Primers targeting ribosomal protein L30 were used as control for ChIP-qPCR analysis with acetyl-H3K27 antibody. Control, Ski, SnoN and Ark represent the expressed sgRNAs. DKO, double knock out. TKO, triple knockout.

---

**Table I.**
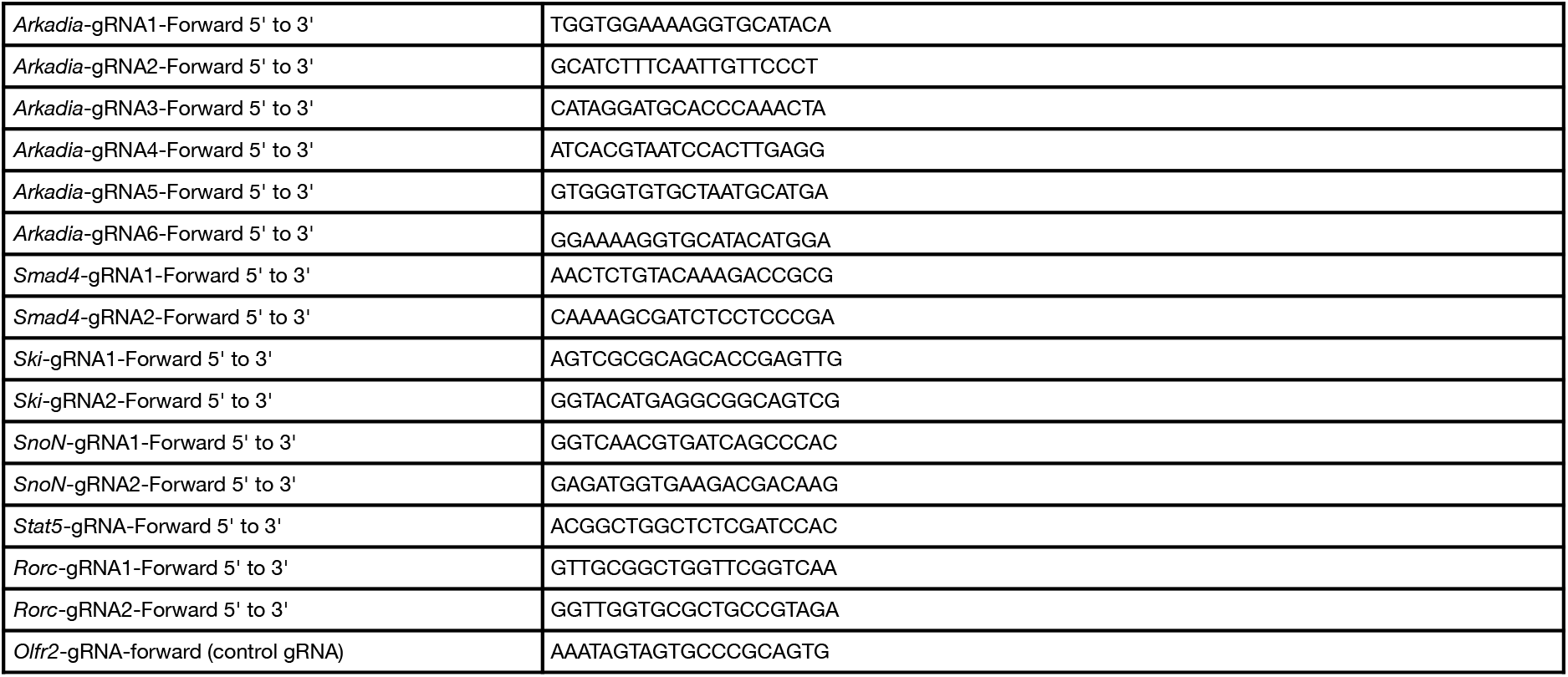
gRNA sequence

**Table II.**
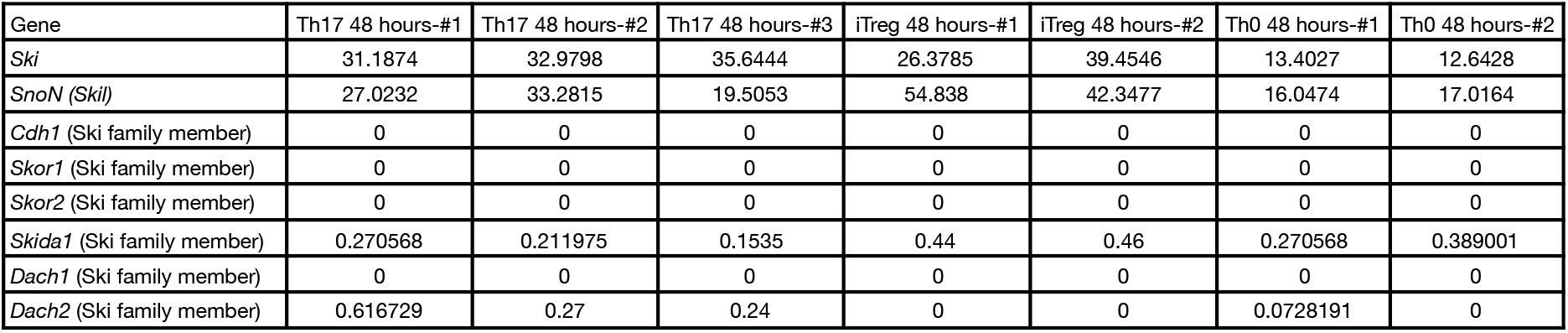
gene expression detected by RNA_seq in cells polarized *in vitro* (FPKM)

**Table III.** antibodies used for cytometry

|  |  |  |
| --- | --- | --- |
| Flow Cytometry: anti-mouse CD4 (RM4-5) AlexaFluor700 | ThermoFisher | Cat. 56-0042 |
| Flow Cytometry: anti-mouse CD4 (RM4-5) eFluor450 | ThermoFisher | Cat. 48-0042 |
| Flow Cytometry: Anti-mouse CD8a (53-6.7), FITC | ThermoFisher | Cat. 11-0081 |
| Flow Cytometry: anti-mouse CD19 (1D3) PerCP-Cy5.5 | TONBO | Cat. 65-0193 |
| Flow Cytometry: anti-mouse CD25 (PC61.5) PE | ThermoFisher | Cat. 12-0251 |
| Flow Cytometry: anti-mouse CD44 (IM7) BV510 | BD Bioscience | Cat. 563114 |
| Flow Cytometry: anti-mouse CD45.1 (A20) PerCP-cy5.5 | BD Bioscience | Cat. 560580 |
| Flow Cytometry: anti-mouse CD45.2 (104) APC | ThermoFisher | Cat. 17-0454 |
| Flow Cytometry: anti-mouse CD62L (MEL-14) APC | ThermoFisher | Cat. 17-0621 |
| Flow Cytometry: anti-mouse TCRβ (H57-597) BV711 | BD Bioscience | Cat. 563135 |
| Flow Cytometry: anti-mouse TCR Vβ6 (RR4-7) PE | Biolegend | Cat. 140004 |
| Flow Cytometry: anti-mouse FOXP3 (FJK-16s) PE-Cy7 | ThermoFisher | Cat. 25-5773 |
| Flow Cytometry: anti-mouse Gata-3 (TWAJ) Alexa Fluor 488 | ThermoFisher | Cat. 53-9966 |
| Flow Cytometry: anti-mouse Helios-3 (22F6) PE | Biolegend | Cat. 137216 |
| Flow Cytometry: anti-mouse RORgt (B2D) APC | ThermoFisher | Cat. 12-6981 |
| Flow Cytometry: anti-mouse RORgt (B2D) PE-eFluor 610 | ThermoFisher | Cat. 61-6981 |
| Flow Cytometry: anti-mouse RORgt (Q31-378) BV421 | BD Bioscience | Cat. 562894 |
| Flow Cytometry: anti-mouse T-bet (O4-46) PE-CF594 | BD Bioscience | Cat. 562467 |
| Flow Cytometry: anti-mouse IL-17A (eBio17B7) Alexa Fluor 488 | ThermoFisher | Cat. 53-7177 |
| Flow Cytometry: anti-mouse IFNγ (XM61.2) PE-Cy7 | ThermoFisher | Cat. 25-7311 |
| Flow Cytometry: anti-mouse IFNγ (XM61.2) PerCP-Cy5.5 | ThermoFisher | Cat. 45-7311 |

**Table IV.**
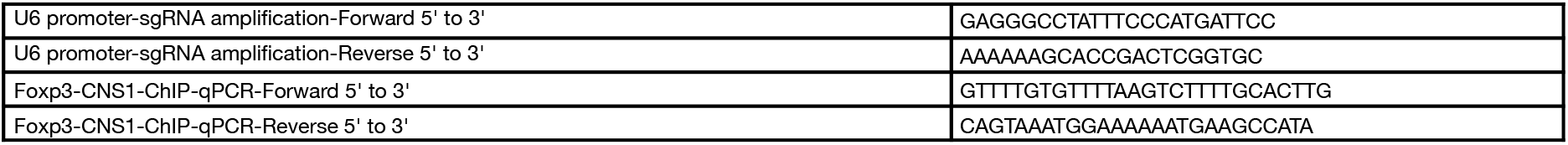
Primers for PCR and ChIP-qPCR

